# A hyperactive transcriptional state marks genome reactivation at the mitosis-G1 transition

**DOI:** 10.1101/053678

**Authors:** Chris C.-S. Hsiung, Caroline Bartman, Peng Huang, Paul Ginart, Aaron J. Stonestrom, Cheryl A. Keller, Carolyne Face, Kristen S. Jahn, Perry Evans, Laavanya Sankaranarayanan, Belinda Giardine, Ross C. Hardison, Arjun Raj, Gerd A. Blobel

## Abstract

During mitosis, RNA polymerase II (Pol II) and many transcription factors dissociate from chromatin, and transcription ceases globally. Transcription is known to restart in bulk by telophase, but whether de novo transcription at the mitosis-G1 transition is in any way distinct from later in interphase remains unknown. We tracked Pol II occupancy genome-wide in mammalian cells progressing from mitosis through late G1. Unexpectedly, during the earliest rounds of transcription at the mitosis-G1 transition, ~50% of active genes and distal enhancers exhibit a spike in transcription, exceeding levels observed later in G1 phase. Enhancer-promoter chromatin contacts are depleted during mitosis and restored rapidly upon G1 entry, but do not spike. Of the chromatin-associated features examined, histone H3 lysine 27 acetylation levels at individual loci in mitosis best predict the mitosis-G1 transcriptional spike. Single-molecule RNA imaging supports that the mitosis-G1 transcriptional spike can constitute the maximum transcriptional activity per DNA copy throughout the cell division cycle. The transcriptional spike occurs heterogeneously and propagates to cell-to-cell differences in mature mRNA expression. Our results raise the possibility that passage through the mitosis-G1 transition might predispose cells to diverge in gene expression states.

## Introduction

Mitosis is accompanied by a dramatic interruption of nuclear processes. In metazoans, the nucleus is disassembled and bulk RNA synthesis ceases (Prescott and Bender 1962). RNA polymerase II (Pol II) and other components of the eukaryotic transcriptional machinery dissociate from chromatin (Gottesfeld and Forbes 1997; Prasanth et al. 2003; Akoulitchev and Reinberg 1998), in part due to mitosis-specific post-translational modifications (Gottesfeld and Forbes 1997; Rizkallah, Alexander, and Hurt 2011). By late telophase, Pol II is known to re-enter the newborn nuclei in bulk and restore global RNA synthesis (Prasanth et al. 2003). However, we lack general principles of how individual genes re-activate transcription at the mitosis-G1 transition.

Many other interphase nuclear processes are also altered globally to varying extents during mitosis. Studies have described such alterations for the recruitment of transcriptional regulators (Raff, Kellum, and Alberts 1994; Martínez-Balbás et al. 1995; A. Dey et al. 2000; Christova and Oelgeschläger 2001; Kruhlak et al. 2001; Zaidi et al. 2003; Young et al. 2007; Z. Yang, He, and Zhou 2008; Blobel et al. 2009; Kadauke et al. 2012; J. Yang et al. 2013; Caravaca et al. 2013; Poleshko et al. 2013; Lake et al. 2014; Lodhi, Kossenkov, and Tulin 2014), deposition of histone variants and modifications (Kruhlak et al. 2001; Kelly et al. 2010; Varier et al. 2010; Wang and Higgins 2012), chromatin structure (Kuo, Iyer, and Schwarz 1982; Michelotti, Sanford, and Levens 1997; Kelly et al. 2010; Kadauke et al. 2012; Hsiung et al. 2014), long-range genome folding (Naumova et al. 2013; Dileep et al. 2015), lamina-associated genomic domains (Kind et al. 2013), and chromosome territories (Walter 2003). Details related to the kinetics, order, and fidelity with which such structures and processes are re-established during the mitosis-G1 transition are largely unknown, except a few examples for factor localization (Prasanth et al. 2003; Poleshko et al. 2013), lamina-associated domains (Kind et al. 2013), and long-range chromosome interactions (Dileep et al. 2015).

Given these uncertainties in the gene regulatory milieu at the mitosis-G1 transition, might there be altered transcriptional output during this cell cycle phase? A microarray-based study identified approximately 200 mature mRNAs that fluctuate during early G1 in mammalian cells (Beyrouthy et al. 2008), but it is unknown to what extent changes in transcriptional activity, versus post-transcriptional modulation, are responsible for these fluctuations. Several studies have directly quantified transcriptional activity over time in cells transitioning from mitosis to interphase (Blobel et al. 2009; Anup Dey et al. 2009; Muramoto et al. 2010; Zhao et al. 2011; Kadauke et al. 2012; Fukuoka et al. 2012; Caravaca et al. 2013), using RT-qPCR of primary transcripts of candidate genes (Blobel et al. 2009; Anup Dey et al. 2009; Kadauke et al. 2012; Fukuoka et al. 2012; Caravaca et al. 2013), live-cell imaging of transcription of *act-5* in Dictyostelium (Muramoto et al. 2010) and a multi-copy reporter locus in a human cell line (Zhao et al. 2011), and microarray-based measurements of nascent transcripts (Fukuoka et al. 2012). Several of these studies suggest or assume that transcriptional output early after mitosis starts off low and rises monotonically with G1 progression at varying kinetics (Blobel et al. 2009; Zhao et al. 2011; Kadauke et al. 2012; Fukuoka et al. 2012; Caravaca et al. 2013). However, some genes show non-monotonic changes in transcriptional output with cell cycle progression after mitosis, but no explanations for these observations have been proposed (Anup Dey et al. 2009; Muramoto et al. 2010; Fukuoka et al. 2012; Caravaca et al. 2013). It remains unclear which transcriptional pattern represents the general rule, as these previous approaches lacked genome-wide extraction of the most prominent patterns. Moreover, some of these studies are difficult to compare due to incongruencies in their temporal coverage of transcriptional measurements, and did not define a clear time frame for the occurrence of the first transcriptional cycle at the mitosis-G1 transition. Major questions remain unresolved: Genome-wide, when does *de novo* transcription upon reversal of mitotic silencing occur? Does the transcriptional program immediately after mitosis deviate significantly from later in interphase, and how might the mitosis-G1 transition influence the fidelity of transcriptional control?

To address these questions, we quantified transcriptional activity from mitosis through G1 phase using three independent methods: chromatin immunoprecipitation-sequencing (ChIP-seq) of Pol II, RT-qPCR of primary transcripts, and simultaneous imaging of nascent and mature mRNA in single cells by single-molecule RNA fluorescence *in situ* hybridization (FISH). The temporal and genomic resolution of our strategy enabled visualization of the pioneering round of transcription at many genes upon reversal of mitotic silencing. We found that during the earliest rounds of transcription most active genes and intergenic enhancers are transcribed at a higher level than later in G1. This observation counters the prevailing assumption of generally lower initial transcriptional outputs immediately after reversal of mitotic silencing. Notably, the mitosis-G1 transcriptional spike does not scale with the frequency of enhancer-promoter chromatin contacts, but is correlated with and preceded by higher levels of histone H3 lysine 27 acetylation (H3K27Ac) in mitosis. Single-molecule RNA FISH demonstrates that the early G1 transcriptional spike can constitute the maximum transcriptional activity in the entire cell cycle, and can propagate to cell-to-cell heterogeneity in mature mRNA levels. We discuss potential contributions of the mitosis-G1 spike in transcriptional compensation for changes in DNA copy number in the cell division cycle, and as a source of gene expression heterogeneity.

## Results

### Pol II ChIP-seq on synchronized and purified cell populations reveals the pioneering round of gene transcription at the mitosis-G1 transition

We performed Pol II ChIP-seq during mitotic exit in murine erythroblast cells, G1E, that lack the hematopoietic transcription factor GATA1 (Weiss, Yu, and Orkin 1997). We used a well-characterized sub-line (G1E GATA1-ER) that expresses a GATA1-estrogen receptor fusion protein, enabling study of transcriptional control in the context of estradiol-inducible gene activation and repression (Weiss, Yu, and Orkin 1997). Tracking Pol II occupancy by ChIP-seq during brief cell cycle phases requires isolating a large number of cells specifically from the desired stages (Fig. 1A). To accomplish this, we arrested G1E GATA1-ER cells (induced with estradiol for 13h) in prometaphase by nocodazole treatment, followed by release into nocodazole-free media for 40min-360min. To minimize contamination with cells from undesired stages of the cell cycle, we purified cells from specific cell cycle phases at specified time points using a fluorescence-activated cell sorting (FACS) strategy (Fig. 1A). This approach is based on a reporter (Kadauke et al. 2012) that consists of YFP fused to a mitotic degradation domain (MD), which confers degradation at the metaphase-anaphase transition (Glotzer, Murray, and Kirschner 1991; Holloway et al. 1993) (Live-cell fluorescence microscopy in Supplemental Video). The combination of synchronization coupled with fluorescence-activated cell sorting (FACS) based on YFP-MD and DNA content enabled isolation of populations highly enriched for cells in prometaphase, between anaphase and cytokinesis, early G1, and late G1 (Fig. 1A). One critical benefit of this strategy is that the G1 samples (sorted for 2N DNA content) are devoid of residual mitotic cells (4N) that might be delayed in their release from nocodazole arrest. Such contamination with transcriptionally silent mitotic cells would lead to an underestimate of the early G1 transcriptional activity in an ensemble assay.

We used these synchronized and sorted populations for ChIP-seq of total Pol II in three biological replicates. Examination of individual loci showed that Pol II ChIP signal is eliminated in prometaphase (Fig. 1B), with minimal residual signal attributable to contamination of this particular sample by ~10% G2-phase cells (Fig. S1), which are also 4N, high YFP-MD. This contamination of the 0h sample does not affect the subsequent time points in our FACS purification strategy (Fig. 1A). Our approach enabled capturing the pioneering round of transcription, which is apparent as a synchronous wave of 5’ to 3’ Pol II progression that initiates between anaphase and cytokinesis (4N, low YFP-MD) at 40 minutes after release (Fig. 1B). This leading edge of Pol II ChIP signal represents a population-averaged position of the first polymerases to travel down a given gene over time, reaching the 3’ end of genes at time points consistent with gene lengths, as shown for illustrative loci in Fig. 1B. The partial progression of the Pol II leading edge can be seen for genes >50kb at individual loci (Fig. 1B) and as a Pol II binding profile averaged across all such genes (Fig. S2). Shorter genes appear to have already completed the first transcriptional cycle, or the first several cycles, some time between the 40min and 60min time points. Thus, the onset of transcriptional reactivation occurs within a narrow window between anaphase and cytokinesis (40min to 60min after release from nocodazole arrest).

**Fig. 1:**
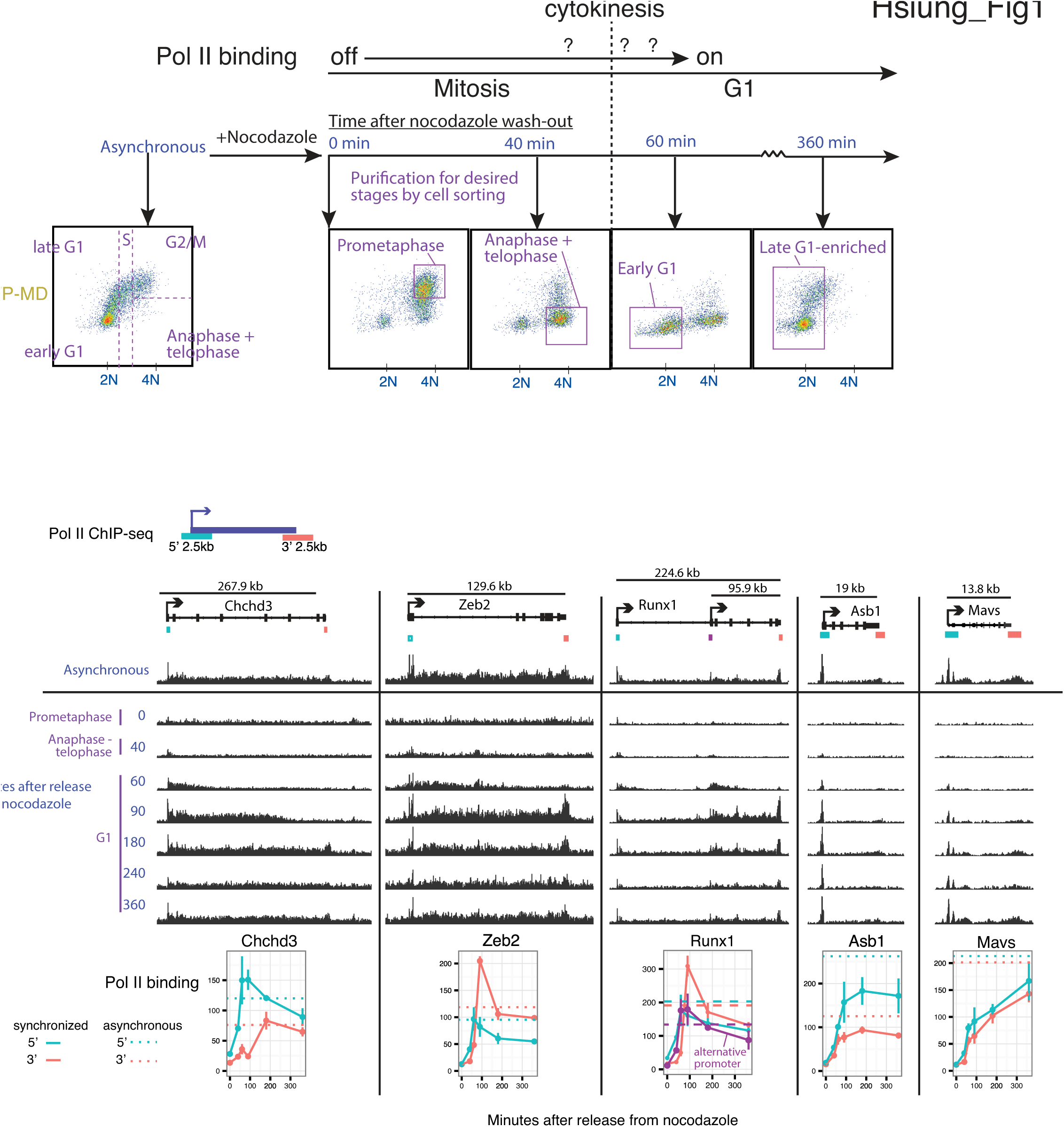
Cell cycle synchronization and purification enables visualizing the pioneering round of transcription at the mitosis-G1 transition by Pol II ChIP-seq. A) Schematic of an experimental strategy that combines nocodazole arrest-release with fluorescence-activated cell sorting (FACS) on the cell cycle reporter YFP-MD (degraded at the metaphase-anaphase transition) and DAPI signals to obtain pure populations from desired cell cycle stages spanning prometaphase through late G1. The subpopulations sorted are demarcated by purple boxes. This strategy ensures that the sorted early G1 sample is devoid of contaminating mitotic cells that are delayed in their release from nocodazole arrest, which could cause an underestimate of transcriptional activity when measured in bulk. B) Sorted cell populations from A) were used for ChIP-seq of total Pol II in biological triplicates, and reads were pooled across replicates. Shown are genome browser track views at illustrative loci to highlight the 5’-3’ progression of the pioneering round of transcription. Y-axes for browser tracks are normalized by library size to enable comparison across time points for each locus, but the y-axes across loci are not meant to be compared in this view. Below the browser tracks, we quantify mean Pol II binding across the 3 replicates over the time course for the 2.5kb regions at the 5’ and 3’ ends of each gene, with error bars indicating SEM. Quantification is also shown for the internal promoter of *Runx1*. All quantifications of Pol II binding in this study are based on library size-normalized read densities (reads per kilobases per million total reads, RPKM).

In addition to the progression of Pol II along the gene body, the amount of Pol II initiating transcription changes in gene-specific patterns over time. For example, at *Chchd3*, *Zeb2* and *Runx1*, Pol II occupancy reaches maximum at the 60-90 min time points at the 5’ region of these genes, followed by a decline through the remainder of G1 phase (Fig. 1B). We refer specifically to this pattern of a sharp increase—followed by sustained decrease—as a “spike.” Importantly, at genes with this particular pattern, the increase in Pol II binding at the 5’ region propagates through the full gene length, visible as a spike in occupancy at the 3’ region with a time delay consistent with gene length. The downward sloping part of the spike indicates that this spike in activity is diminished shortly after the completion of the initial transcriptional cycles. The temporal spike in Pol II binding at the 3’ end that follows that at the 5’ end in time (exemplified by Chchd3, Zeb2, and Runx1 in Fig. 1B) indicates that the spike in Pol II binding reflects full-length transcription of the gene, rather than just an increase of paused Pol II at the 5’ end. Not all genes display a transcriptional spike; for example, at *Asb1*, Pol II binding plateaus after ~90 min of release (Fig. 1B), whereas at *Mavs*, Pol II binding rises continuously over a period of 360min following release (Fig. 1B).

### A spike in transcriptional activity at the mitosis-G1 transition is prevalent across the genome

To examine global distributions of Pol II occupancy over these time points, we measured Pol II occupancy at the 5’ regions of the 4309 non-overlapping genes with above-background binding in at least one time point, as determined by a peak-caller (Zhang et al. 2008). Globally, Pol II binding reaches substantial levels above background even prior to the completion of the first round of transcription for many genes at 60min after release (Fig. 2A). In terms of the rise in absolute Pol II binding prior to the 60min time point, the onset of transcription occurs globally with minimal gene-to-gene differences in kinetics within the limits of our temporal resolution. Pol II binding at the 60min and 90min time points overall overshoot that of the 360min (Fig. 2A; individual replicates shown in Fig. S5). By 240min, the distribution returns to roughly the same as 360min (Fig. 2A). Thus, contrary to prior expectations of transcriptional reactivation post-mitosis starting off with generally lower initial output, transient transcriptional hyperactivity is a widespread phenomenon associated with the earliest rounds of transcription upon reversal of mitotic silencing.

**Fig. 2:**
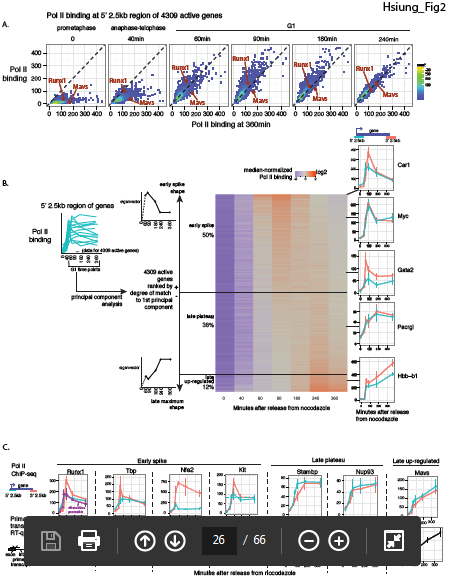
A spike in gene transcription is prevalent across the genome at the mitosis-G1 transition. A) Replicate-averaged Pol II binding at the 5’ 2.5kb region of 4309 genes active in at least one time point is plotted for each time point against the 360min time point. *Runx1* and *Mavs*, two genes with distinct temporal patterns shown in Fig. 1B, are highlighted. Plots for individual replicates are shown in Fig. S5B. B) For the same 4309 genes as in A), we performed principal component analysis on Pol II binding (RPKM), normalized by total Pol II binding for each gene over all time points, at the 5’ 2.5kb region of each gene. For this analysis, only G1 time points (60min, 90min, 180min, 240min, 360min) were used. The temporal “shapes” of Pol II binding (eigenvector) of the first principal component is shown. Genes were ranked by their degree of match to (projection onto) the first principal component, and all their gene-normalized RPKM at the 5’ 2.5kb regions plotted in a heatmap for all time points. Threshold for “early spike” is defined as the inflection of projection onto the first principal component from positive to negative. The threshold for separating “late plateau" and “late up-regulated" were chosen manually based on the appearance of the heatmap. 434 early spike genes and 432 late-upregulation genes meet significance threshold (p < 0.05, determined by bootstrapping) for their projections onto the first principal component (Fig. S5A). The positions of several genes in the heatmap are shown on the right, together with their RPKMs for both the 5’ 2.5kb and 3’ 2.5kb regions. C) Pol II ChIP-seq binding profiles (error bars denote SEM, n = 3) are shown together with quantification of primary transcripts by RT-qPCR using primers flanking intron-exon junction (error bars denote SEM, n=5-6). Pol II ChIP profiles from Runx1 and Mavs from Fig. 1B are reproduced here for ease of comparison. In addition to these genes, profiles for Gata2, Kit, Hbb-b1, Hba-a1 are shown in A) for Pol II ChIP and in Fig. 3D for primary transcript RT-qPCR.

In general, comparisons of factor occupancy across ChIP-seq samples in the context of global changes in binding require that changes in normalized read counts accurately reflect absolute changes in binding. This important property holds true in our data due to the presence of a relatively large proportion of reads mapping to intergenic regions that represent non-specific background (Fig. S3). This background serves as an internal calibration across sequencing libraries, enabling inferences of changes in Pol II occupancy on an absolute scale (Fig. S3). We also confirmed patterns observed by Pol II ChIP-seq at individual loci by Pol II ChIP-qPCR (Fig. S4), further indicating that sequencing read counts reflect quantitation by qPCR.

While transcriptional hyperactivity upon reversal of mitotic silencing is a prevalent trend, individual genes can exhibit a variety of distinct temporal profiles of transcription, indicating a degree of gene specificity for such patterns (Fig. 1B). To stratify Pol II binding patterns at individual genes in an unbiased manner, we performed principal component analysis on Pol II binding at the 5’ region of genes at G1 phase time points (60min-360min). For this analysis, we first normalized Pol II binding at each time point by the sum of Pol II binding across all time points to remove gene-to-gene differences in transcriptional activity unrelated to cell cycle progression. The first principal component accounts for the most (47.2%) gene-to-gene variance and represents temporal shapes that fall along a continuum of early G1 spike vs. late G1 up-regulation in Pol II binding (Fig. 2B). The temporal shapes of individual genes, as defined by the projection onto the first principal component, is highly concordant across the three biological replicates (R = 0.8-0.9, Fig. S6). Lower ranking principal components are less clearly distinguishable from noise (Fig. S6). Ranking genes based on the degree of match to the first principal component (projection of each gene onto this principal component) reveals that approximately 50% of genes exhibit an early spike, 38% late plateau, and 12% late up-regulation in Pol II binding, though these patterns are not discrete clusters (Fig. 2B). Among the 50% of genes exhibiting some degree of match to the early spike pattern, the magnitude of the spike at the 90min time point is on average 1.4-fold, and can reach up to 4.3-fold, higher than the 360min time point (Fig. S7). Hereafter we refer to the early G1 spike as a trait defined quantitatively by the degree of match in the positive orientation of the first principal component as shown in Fig. 2 xsB.

We found no association between the early G1 spike and the traveling ratio of Pol II, indicating that the occurrence of the early transcriptional spike does not involve a difference in the rate of Pol II promoter escape (Fig. S8). Pol II binding at the 3’ regions of genes often mirrors the temporal shape for the corresponding 5’ regions (Fig. 2B panels on right, and Fig. 2C top row), and very similar principal components were obtained from applying the analysis to Pol II binding at the 3’ region of genes (analysis not shown). Thus, these temporal changes in Pol II binding reflect full-length gene transcription. Indeed, RT-qPCR of primary transcripts using primers flanking intron-exon junctions for a subset of genes demonstrate that the temporal patterns of Pol II binding at individual loci are well reflected at the level of RNA synthesis (Fig. 2C bottom row).

The early G1 transcriptional spike pattern encompasses genes with functions general to many cell types (e.g. Tbp, Fig. 2C), as well as genes involved in developmental regulation relevant to hematopoietic cells, such as *Gata2*, *Myc*, *Kit*, and *Runx1* (Fig. 2B and Fig. 2C), with an enrichment for genes in p53 signaling pathways (Fig. S9). The late up-regulation pattern enriches for Gene Ontology terms related to plasma membrane proteins (Fig. S9), and of relevance to erythroid biology, includes both alpha-globin (*Hba-a1*) and beta-globin (*Hbb-b1*) genes. Note that the late up-regulation category does not necessarily represent delayed transcriptional reactivation on an absolute level; rather, these include genes that tend to reach similarly high levels of Pol II binding at the 60min-90min time points, then exhibit further sustained up-regulation through the remainder of G1, as exemplified by absolute Pol II binding for *Mavs* (Fig. 1B) and *Hbb-b1* (Fig. 2B).

To test whether the early G1 spike versus late G1 up-regulation patterns are cell-type specific, we performed nocodazole-mediated mitotic arrest-release in a murine embryonal carcinoma cell line (F9) (ALONSO’ et al. 1991) and measured transcription by primary transcript RT-qPCR. Of 13 genes examined that are expressed in both G1E GATA1-ER and F9 cells, 10 showed similar G1 transcriptional patterns in both cell types, indicating that the G1-phase modulation of transcription we uncovered in G1E GATA1-ER cells can be found across developmentally distinct murine cell types (Fig. S10).

### Mitosis-G1 transcriptional spike also occurs at intergenic enhancers, but restoration of enhancer-promoter chromatin contacts after mitosis does not spike

Pol II is known to transcribe not only genes, but also enhancers. Do enhancers display similar transcriptional outputs as that of genes at the mitosis-G1 transition? We previously identified enhancers in G1E GATA1-ER cells based largely on DNase-sensitive sites that do not overlap known transcriptional start sites, and coincide with presence of H3K4me1 in the relative absence of H3K4me3 (Hsiung et al. 2014). We quantified the level of Pol II binding at enhancers in the mitosis-G1 time course of G1E GATA1-ER cells from this study. We restricted our analysis to a set of 809 active enhancers with above-background Pol II binding and located away from genes (>3kb from 5’ end and >10kb from 3’ end of gene annotations) to avoid confusion with signal arising from Pol II occupancy at genes, which can extend several kilobases beyond the 3’ end of the gene. Principal component analysis performed on Pol II binding at these intergenic enhancers showed that the top principal component (Fig. 3A; replicate concordance analysis in Fig. S11, R = 0.78-0.88) reflects temporal shapes similar to that obtained from the analysis of genes in Fig. 2B. Analogous to our analysis for genes in Fig. 2B, we stratified Pol II binding patterns at enhancers into early spiking, late plateau, and late up-regulated patterns based on the degree of match to the first principal component (Fig. 3A). Approximately half of all examined enhancers display an early spike in Pol II occupancy.

**Fig. 3:**
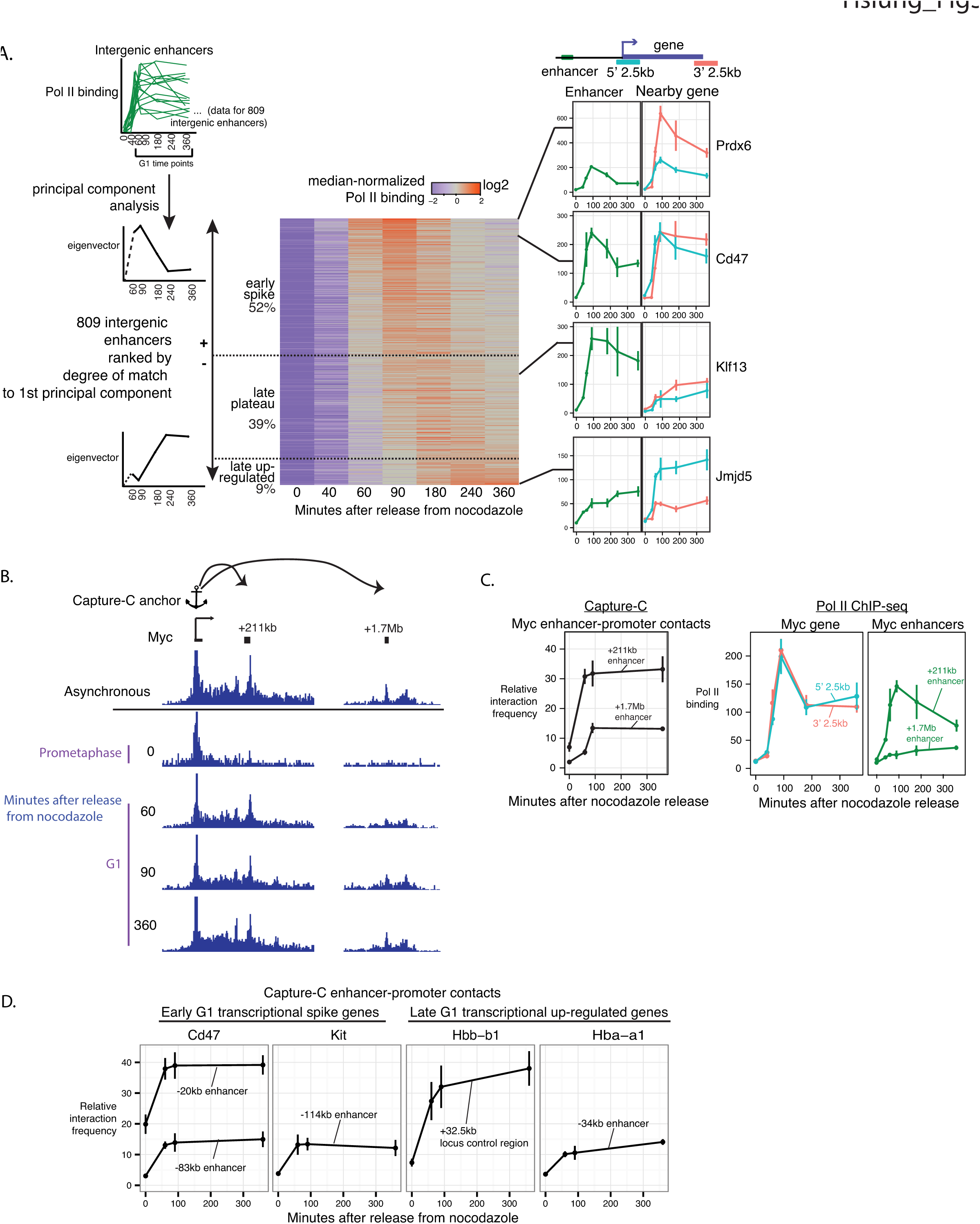
Pol II binding at enhancers, but not enhancer-promoter contacts, also spikes at the mitosis-G1 transition. A) For 809 intergenic enhancers, we performed principal component analysis on Pol II binding in the same fashion as that detailed for genes in Fig. 2B. Shown are results outlined in a fashion analogous to Fig. 2B. Additionally, for the enhancers we highlight to the right of the heatmap, we also show the raw RPKM of Pol II binding at the nearest gene. B) Capture-C in a nocodazole arrest-release using the Myc promoter as the anchor. Y-axis of browser tracks are read counts of ligation products normalized to total number of ligation products in the library. We highlight a likely enhancer at the +211kb region (resides within transcribed region of non-coding RNA Pvt1, omitted in graph for clarity) and a known enhancer at the +1.7Mb region (Shi et al. 2013). C) Left: Quantification of Capture-C read densities for the Myc +211kb and +1.7Mb enhancers. Y-axis denotes read counts reflecting ligation products between enhancer region and the anchor, normalized to total number of ligation products in the library. Error bars denote SEM, with n = 3 sequencing libraries encompassing 2 separate ligation libraries and 3 separate oligo captures. Right: Pol II ChIP-seq read densities at the Myc gene (5’ 2.5kb and 3’ 2.5kb, duplicated from Fig. 2 for ease of comparison) and at the +211kb and +1.7Mb enhancers. D) Quantification of enhancer-promoter contacts using Capture-C anchors at the promoters of early G1 transcriptional spike genes (Cd47, Kit) and late G1 upregulated genes (Hba-a1, Hbb-b1). See Fig. S14 for browser tracks. These enhancers were described in prior studies: Cd47 −20kb (Dogan et al. 2015), Kit −114kb (Jing et al. 2008; Lee et al. 2015), Hba-a1 −34kb R2 region (Hughes et al. 2005), and Hbb-b1 +32.5kb locus control region (Bender et al. 2000). Y-axis is normalized Capture-C contact frequency as described in C). Error bars and number of replicates are as described in C), except for Hbb-b1, which represents n = 2 of separate ligation libraries.

How do Pol II binding patterns at enhancers relate to those of nearby genes? At some loci, such as the Cd47 locus, the enhancer and its nearest gene exhibit the early spike pattern (Fig 3A). However, some loci exhibit a transcriptional spike for the gene without appreciable spiking of Pol II binding at its known enhancer (Fig. 3A and Fig. S12A). When examined in an unbiased manner across the 809 intergenic enhancers, the correlation with the degree of early G1 Pol II spiking at the nearest gene is mild (R=0.34, Fig. S13) and hence difficult to interpret. We note that assigning each enhancer to its nearest gene does not account for enhancers that regulate distant genes, and that there is not necessarily a one-to-one pairing of enhancers and genes.

As another proxy for enhancer activity, we measured enhancer-promoter chromatin contacts using Capture-C (Hughes et al. 2014), a multiplexed derivative of chromosome conformation capture, in a mitotic arrest-release time course with anchors located at promoters of 3 early spike genes (Cd47, Kit, Myc) and 2 late up-regulation genes (Hba-a1, Hbb-b1). Enhancers are known to preferentially lose chromatin accessibility during mitosis (Hsiung et al. 2014), but it is unknown whether enhancer-promoter chromatin contacts can be maintained during mitosis. We found that all of the enhancer-promoter pairs examined showed depletion of contacts during mitosis (Fig. 3B-D, Fig. S12B, and Fig. S14). These results demonstrate that mitotic disruption of long-range genome folding—previously shown for replication timing domains (Dileep et al. 2015), and chromosome compartments and topologically-associating domains (Naumova et al. 2013)—is a general rule that includes enhancer-promoter contacts.

Upon G1 entry, the enhancer-promoter contacts increase sharply by 60min-90min and show no significant change by 360min, regardless of whether the gene transcriptional pattern is categorized as early spike or late up-regulated (Fig. 3B-D). We conclude that the frequency of enhancer-promoter contacts do not necessarily scale quantitatively with early G1 spike or late G1-upregulation of transcription. Whether enhancer-promoter contacts are required to initiate early G1 transcriptional spike at genes remains an open question.

### Mitotic H3K27Ac levels predict mitosis-G1 transcriptional spiking at genes and intergenic enhancers

What mechanism might underlie the mitosis-G1 transcriptional spike at genes and enhancers? We hypothesized that chromatin-associated features, especially those during mitosis, at genes and enhancers may predict differences among loci in their G1 transcriptional patterns. We performed correlative analysis of the following data: 1) Pol II ChIP-seq in asynchronous cells (data generated in this study); 2) DNase-seq in mitotic and asynchronous cells (from (Hsiung et al. 2014)), 3) transcription factor GATA1 ChIP-seq in mitotic and asynchronous cells (from (Kadauke et al. 2012)); 4) histone H3 lysine 27 acetylation (H3K27Ac) ChIP-seq in mitotic and asynchronous cells (data generated in this study); and 5) ChIP-seq of histone H3 lysine methylation modifications in asynchronous cells (H3K4me3, H3K4me1, H3K27me3 and H3K9me3 data from (Wu et al. 2011)). Importantly, the mitosis data sets for DNase-seq and GATA1 ChIP-seq were derived from nocodazole-arrested cells subjected to purification by FACS for the mitotic epitope H3S10Ph to achieve >98% mitotic purity (Hsiung et al. 2014; Kadauke et al. 2012). Likewise, to carry out H3K27Ac ChIP-seq on mitotic cells, we applied a similar procedure using a more robust and cost-effective mitosis-specific antibody (MPM2) to achieve essentially 100% mitotic purity (Campbell, Hsiung, and Blobel 2014). The mitotic purity of these samples ensures that the data reflect properties of mitotic cells, rather than contaminating interphase cells. H3K27Ac levels in prometaphase are strongly, but imperfectly, correlate with that in asynchronous cells at promoters (R = 0.72) and intergenic enhancers (R = 0.72), indicating a degree of locus specificity in the maintenance of H3K27Ac in mitosis (Fig. S15). To our knowledge, this is the first report of H3K27Ac levels in mitosis measured by ChIP.

For each data set, we quantified read counts at promoters of active genes (4309 genes analyzed in Fig. 2) and intergenic enhancers (809 enhancers analyzed in Fig. 3). In Fig. 4, we show the Pearson correlation coefficient between the read counts for each feature with our measure of early spike vs. late up-regulation G1 transcriptional patterns (Pol II ChIP-seq degree of match to the first principal component in Fig. 2 and Fig. 3), analyzing promoters and intergenic enhancers separately. Among the features examined at promoters, the early G1 transcriptional spike is most strongly correlated with H3K27Ac levels in mitosis, including both prometaphase (R = 0.47) and anaphase-telophase (R = 0.53) populations. By comparison, the correlation at promoters is less positive with H3K27Ac levels in asynchronous cells (R = 0.30), indicating that mitotic H3K27Ac levels specifically provides some additional predictive power. Likewise, for intergenic enhancers, H3K27Ac level in mitosis (R = 0.35 for prometaphase, R = 0.46 for anaphase-telophase) is the most positively correlated with the early G1 transcriptional spike, and is less positively correlated for H3K27Ac level in asynchronous cells (R = 0.12) (Fig. 4).

**Fig. 4:**
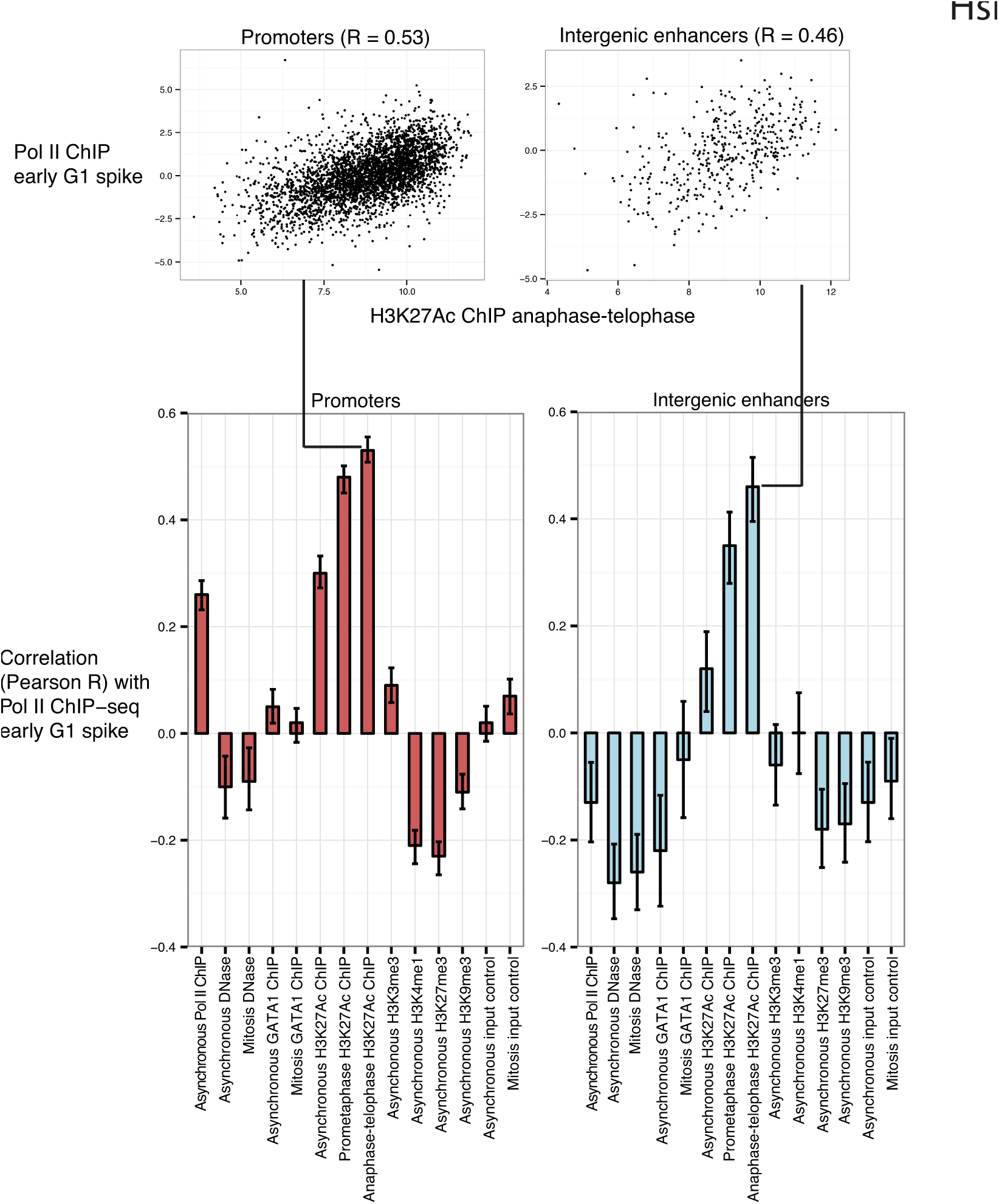
Higher levels of H3K27Ac during mitosis predict the mitosis-G1 transcriptional spike at genes and intergenic enhancers. We examined the signal strength of chromatin-associated features in mitotic and/or asynchronous cells for their genome-wide correlation (Pearson R) with the early G1 transcriptional spike, defined in Fig. 2B for genes and in Fig. 3A for intergenic enhancers as the degree of match to the first principal component from Pol II ChIP-seq. Pol II ChIP-seq generated in this study; DNase-seq from (Hsiung et al. 2014); GATA1 ChIP-seq from (Kadauke et al. 2012); H3K27Ac ChIP-seq generated in this study; H3K4me1, H3K4me3, H3K9me3, H3K27me3 from (Wu et al. 2011). The mitosis H3K27Ac ChIP-seq sample consisted of approximately 100% mitotic purity obtained by FACS for MPM2 positivity as described in (Campbell, Hsiung, and Blobel 2014). Error bars denote 95% confidence intervals.

In contrast, the mitotic or asynchronous levels of two features at promoters previously shown to have locus-specific degrees of persistence in mitosis—DNase sensitivity (Hsiung et al. 2014) and GATA1 occupancy (Kadauke et al. 2012)—showed no correlation with the early G1 transcriptional spike. However, this does not rule out that there might be a minimal level of mitotic DNase sensitivity required for the early G1 transcriptional spike, because the promoters of virtually all transcriptionally active genes are at least somewhat DNase-sensitive in mitosis. The early G1 transcriptional spike is also weakly correlated with promoter levels of Pol II binding (R = 0.26), suggesting a mild association with overall levels of transcriptional activity (Fig. 4). Another active promoter modification, H3K4me3, is not predictive (R = 0.09), whereas levels of repressive modifications H3K27me3 (R = −0.23) and H3K9me (R = −0.11) in asynchronous cells are weakly anti-correlated with the early G1 transcriptional spike (Fig. 4). At intergenic enhancers, levels of DNase sensitivity, GATA1 occupancy, H3K4me1, H3K4me3, H3K27me3, and H3K9me3 in mitotic and/or asynchronous cells are all either weakly anti-correlated or uncorrelated with the early G1 transcriptional spike (Fig. 4).

We conclude that H3K27Ac levels specifically during mitosis exceed other indicators of active chromatin in its predictive power of the early G1 transcriptional spike at both genes and intergenic enhancers. Since this strongest predictor precedes the mitosis-G1 transcriptional spike, the temporality of the association is consistent with the possibility that mitotic H3K27Ac may be involved in causing the mitosis-G1 transcriptional spike.

### Mitosis-G1 transcriptional spike propagates to cell-to-cell heterogeneity in mature mRNA expression

Our findings thus far demonstrate a spike in transcriptional activity at the mitosis-G1 transition based on measurements of cell population average. Does this transcriptional spike occur in all cells in the population or only in a subset of cells, thus potentially contributing to transcriptional heterogeneity among cells? Is the mitosis-G1 transcriptional spike buffered by post-transcriptional regulation, or does the transcriptional spike propagate to elevated mature mRNA levels? To address these questions, we used single-molecule RNA fluorescence in situ hybridization (FISH) to simultaneously quantify nascent and mature mRNA in single cells by 3D microscopy (Femino 1998; Raj et al. 2008; Levesque and Raj 2013). We imaged nascent and mature mRNAs in the same field by hybridizing fixed cells with spectrally distinguishable probes specific to introns or exons of a given gene. While the vast majority of exonic probe signals are from mature mRNAs, co-localized exonic and intronic probe signals are primary transcripts that mark active transcription sites in interphase cells (Fig. 5A). In prometaphase, cells with condensed chromosomes show no detectable signal in the intronic channel due to mitotic transcriptional silencing, whereas stable mature mRNA molecules that presumably arose prior to mitosis are detectable (Fig. 5A). Consistent with our Pol II ChIP-seq data, the earliest active transcription appears in cells between anaphase and cytokinesis (4N, low YFP-MD). This RNA synthesis occurs amidst chromosomes that are still morphologically condensed, demonstrating that overt condensation does not prohibit gene transcription (Fig. 5A and Fig. S16).

**Fig. 5:**
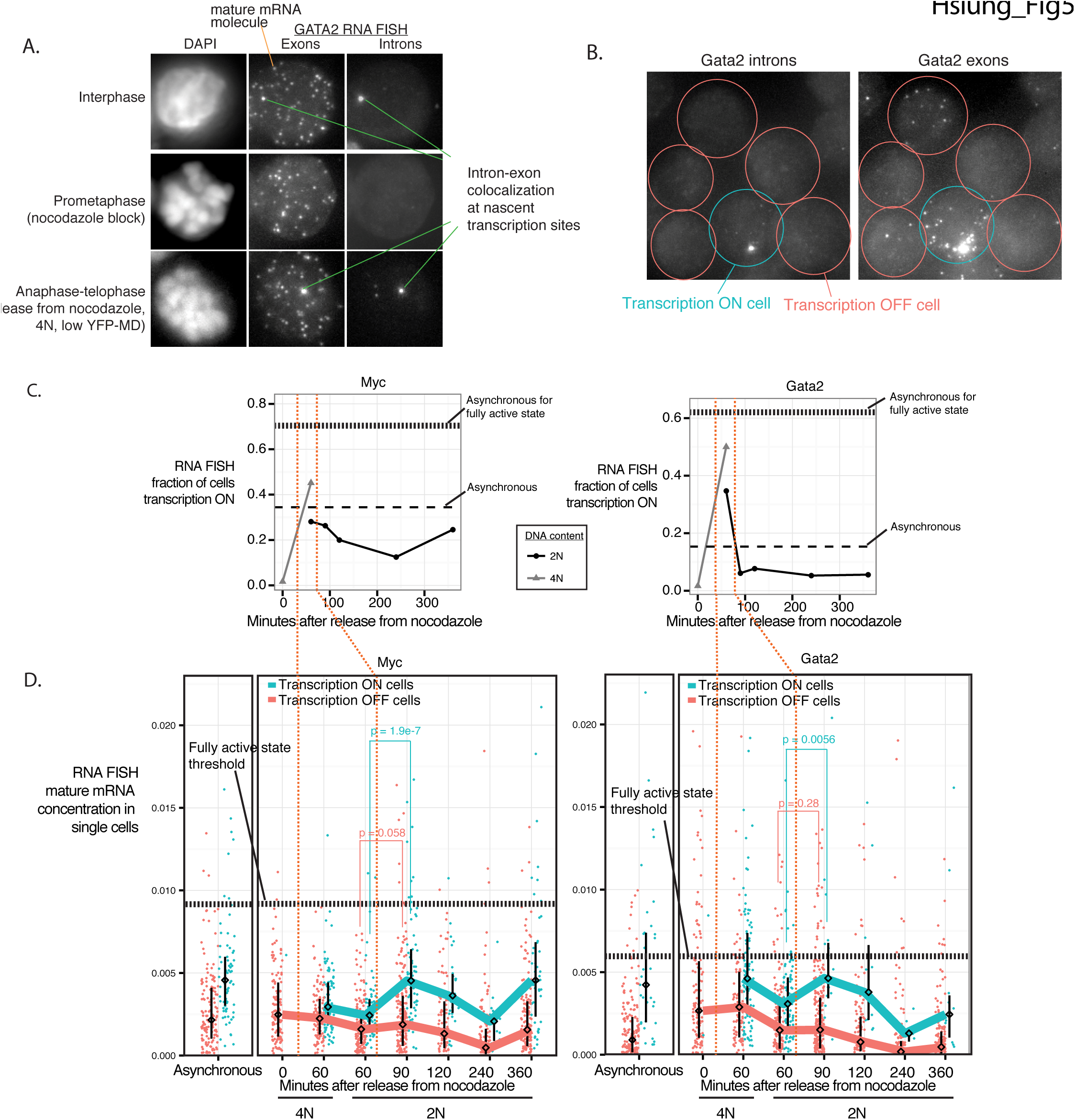
Mitosis-G1 transcriptional spike can propagate to cell-to-cell heterogeneity in mature mRNA expression. A) Images of representative cells in interphase, prometaphase arrest by nocodazole, and between anaphase-telophase (60min after nocodazole release, sorted for 4N and YFP-MD low), taken by 3D wide-field microscopy. Single optical plane is shown for DAPI channel. Maximum projections are shown for Gata2 exonic probe (coupled to Cy3) and intronic probe (coupled to Alexa594) channels for RNA FISH, with single mature mRNA molecules and intron-exon colocalized spots highlighted. B) *Gata2* intron and exon RNA FISH in a representative field of asynchronous cells. C) In cells subjected to nocodazole arrest-release and FACS purified in a manner similar to Fig. 1B, we performed RNA FISH for Myc and Gata2 in the low-expression state (+estradiol 13h), and quantified the fraction of cells that contain at least one intron-exon colocalized spot, indicative of active transcription (referred to as transcriptionally “on”). Horizontal dashed lines indicate levels from asynchronous populations in both the low (+estradiol 13h) and fully activated (no estradiol) states. D) From the same experiment as in C), we show the distributions of single-cell mature mRNA concentration (mRNA count / cell size) of transcriptionally “on” versus “off” cells, with each dot representing a cell. The line connects the medians of the distributions, and the interquartile range is indicated. We imaged 120-262 cells for each time point. Vertical orange dotted lines demarcate timing of transcriptional spike as shown in C). Horizontal dotted lines mark thresholds selected based on receiver operating characteristics curves (Fig. S19) for discriminating the low-expression (+estradiol 13h), versus fully activated (no estradiol), asynchronous populations. Additional biological replicates are shown in Fig. S17. P-values (one-sided Wilcoxon test) for the difference in distributions between the transcription “on” and “off” cells are indicated for select time points.

We focused on two genes that exhibit transcriptional spikes at the mitosis-G1 transition, *Myc* and *Gata2* (Pol II ChIP patterns for both in Fig. 2B). *Myc* and Gata2 encode for transcription factors that regulate stemness and self-renewal, and whose mature mRNA half-lives are relatively short (~15min-2h for Myc, (Dani et al. 1984; Watson 1988; Herrick and Ross 1994)~2.8h half-life for Gata2, (Sharova et al. 2009)). Expression of *Myc* and *Gata2* are down-regulated upon exposure to estradiol for 13h through transcriptional repression by GATA1-ER (Rylski et al. 2003; Grass et al. 2003), and at this relatively low level of expression the mature mRNA levels among single cells can vary by >100-fold (Fig. S19 and Fig. 5C). The single-cell expression patterns of these genes allow evaluation of the degree to which mitosis-G1 transcriptional spiking of these genes can contribute to heterogeneity in mature mRNA levels.

Transcription visualized by single-cell imaging is known to occur intermittently, with intervals of active RNA synthesis interspersed with periods of inactivity (Golding et al. 2005; Chubb et al. 2006; Raj et al. 2006). When cells are fixed and viewed as a static image of single-molecule RNA FISH, this pulsatile nature of transcription manifests as a mix of transcriptionally “on” and “off” cells (Fig. 5B and Fig. S16). Single-molecule RNA FISH shows that the early transcriptional spike we observed as an average across cell populations (Fig. 2B) occurs by a spike in the fraction of cells actively transcribing at the mitosis-G1 transition for both Myc (Fig. 5C left panel; 45% for 4N cells at 60min point vs. 12.5% at 240min time point) and Gata2 (Fig. 5C right panel; 50% for 4N cells at 60min time point vs. 5% at 240min time point). Of note, the zenith of the transcriptional spike coincides with the time point when chromosomes are still morphologically condensed (Fig. 5A and Fig. 5C). In contrast, the intensity of transcription sites are relatively unchanged (Fig. S18). Thus, the mitosis-G1 spike in averaged transcriptional output mostly arises from an increase in the probability of the gene being in a transcriptionally “on” state, not from an increase in the number of nascent transcripts synthesized during each “on” period.

How does the spike in probability of being in a transcriptionally “on” state at the mitosis-G1 transition manifest at the level of mature mRNA in single cells? To address this, we quantified the number of mature mRNA molecules in each cell. For both *Gata2* and *Myc*, the transcriptionally “on” cells express higher levels of mature mRNA than the transcriptionally “off” cells across all time points (Fig. 5D), confirming that the intermittent nature of transcription for these genes contributes visibly to cell-to-cell variability in mature mRNA levels. Furthermore, among the transcriptionally “on” cells, the mature mRNA levels spike at the 90min-120min time points, subsequent to the spike in transcription for both Myc (Fig. 5D left panel; 1.9-fold increase in median mature mRNA concentration from 60min-90min in 2N cells) and Gata2 (Fig. 5D right panel; 1.5-fold increase in median mature mRNA). Of note, this spike is not observed for the transcriptionally “off” cells in the corresponding time points. Additional biological replicates are shown in Fig. S17. In static images, the spike in mature mRNA specifically among transcriptionally “on” cells at the 90min-120min time points (Fig. 5D) must have arisen from transcriptional activity prior in time; thus, the spike in mature mRNA levels at these time points are enriched for the subset of cells that previously participated in the transcriptional spike at the 60min time point. These results support that the mitosis-G1 transcriptional spike reflects an increased probability for individual cells to be in a transcriptionally “on” state, and is substantial enough to produce a spike in mature mRNA levels to overcome any potential buffering by post-transcriptional regulation. While unable to directly offer mRNA expression trajectories of single cells over time, these data suggest that participation by individual cells in the transcriptional spike at the 60min time point may predispose those cells to be in a transcriptionally “on” state in subsequent time points, even when the overall fraction of the population transcribing has already declined.

### Early spike and late-upregulation G1 transcriptional patterns can be observed in the absence of cell cycle synchronization, and can constitute the maximal transcriptional activity per DNA copy in the entire cell cycle

Our analyses thus far have relied on the use of cell cycle synchronization methods, as have previous studies of transcription at the mitosis-G1 transition (Blobel et al. 2009; Anup Dey et al. 2009; Zhao et al. 2011; Fukuoka et al. 2012; Caravaca et al. 2013). While the resolution of cell cycle synchrony provided unambiguous visualization of the pioneering round of transcription upon reversal of mitotic silencing (Fig. 1B), effects of synchronization on transcription are unknown and could potentially confound our gene expression observations. To avoid cell cycle synchronization completely, we sought to measure transcription in cells from different cell cycle stages in an asynchronous population by imaging. We used cell area as a proxy for cell cycle progression since cytokinesis, based on an empirically determined proportionality between the two variables (Fig. S20). We also demonstrated, by tracking live cells through cell divisions using brightfield microscopy, that newly divided cells in early G1 are enriched among the smallest cells (Supplemental Video). Thus, combining RNA FISH with cell area provides a view of transcription with respect to approximate cell cycle progression since cytokinesis, enabling resolution within G1 phase that is difficult to achieve with other cell cycle markers. Satisfyingly, the transcriptional patterns for *Gata2* and *Kit* (early G1 spike genes) and *Hbb-b1* (late G1 up-regulation gene) obtained by this method reflect that measured from approaches using synchronization (Fig. 6). Furthermore, after normalizing for DNA copy number changes, the early G1 spike for *Gata2* and *Kit* constitutes the highest transcriptional activity throughout the cell cycle for those genes, whereas the maximal activity for *Hbb-b1* is near the late G1/S boundary (Fig. 6). Thus, the G1-phase transcriptional patterns we uncovered can be observed in naturally dividing cells in the absence of synchronization, and can constitute periods of maximal activity in the entire cell cycle. We also note that for both early spike genes, Gata2 and Kit, this contributes to a doubling of transcriptional activity per DNA copy when averaged across all of G1, relative to that in G2. In the Discussion, we explore the implications of this for gene dosage compensation for DNA copy changes during the cell division cycle.

**Fig. 6:**
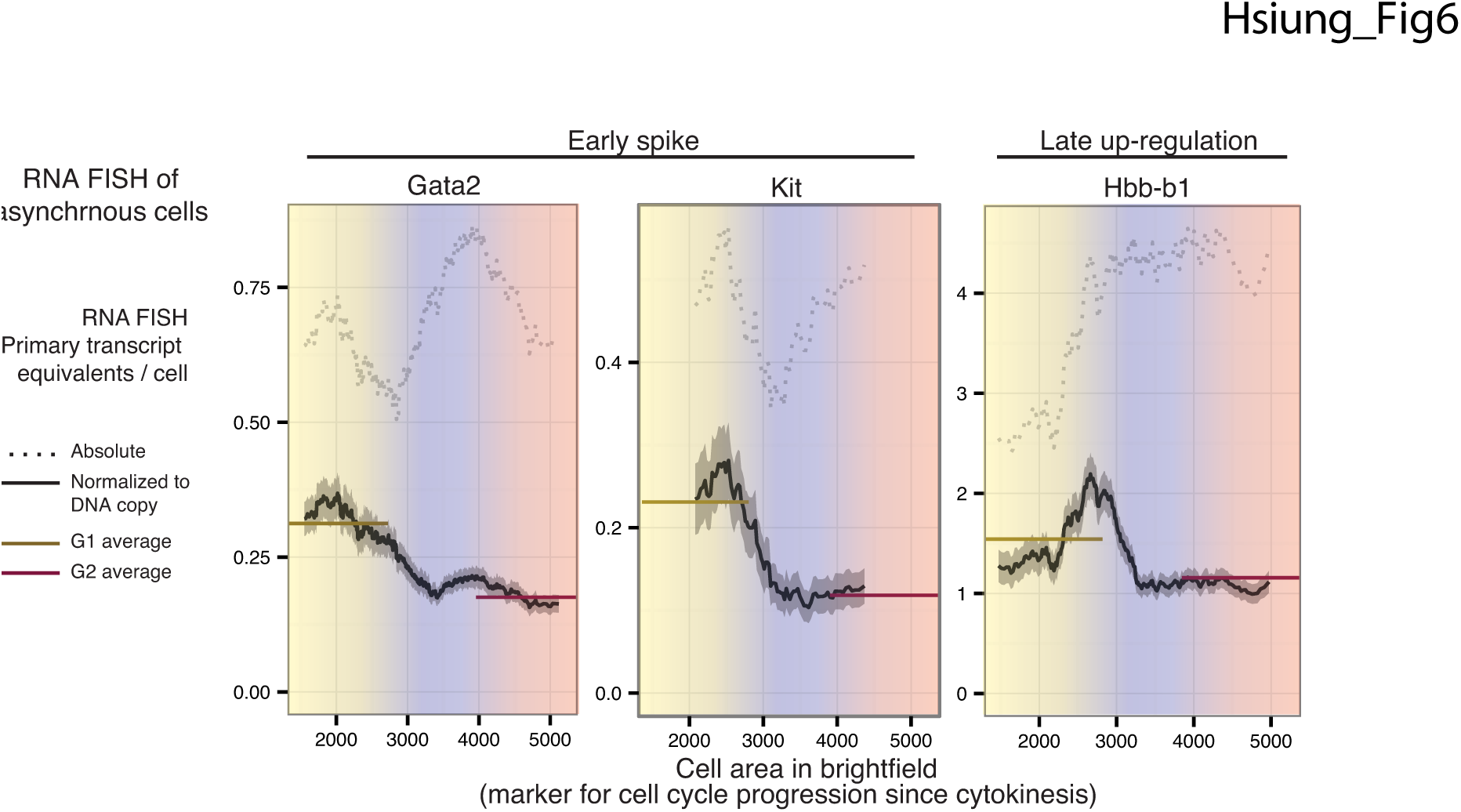
Mitosis-G1 transcriptional spike can be observed in the absence of synchronization and can constitute the maximum transcriptional activity per DNA copy in the entire cell cycle. Asynchronously dividing cells were imaged by 3D wide-field microscopy after performing RNA FISH for two early spike genes (Gata2 and Kit) and a late-upregulated gene (Hbb-b1) using intronic and exonic probes. Primary transcript content is shown in dotted line in terms of absolute primary transcript equivalents per cell, and shown in solid line after normalization by DNA content, as a moving mean across cell area manually determined from brightfield images. Using the proportionality between cell area and DAPI intensity (Fig. S20), we roughly estimate G1, S, and G2-phase boundaries demarcated by color. Solid horizontal lines indicate average within the entire G1 or G2 compartment. Gata2 quantification is based on images of ~5000 cells pooled across 4 biological replicates; Kit quantification is based on images of ~650 cells; Hbb-b1 quantification is based on images of ~1000 cells. Gray shade around moving mean denotes SEM within sliding window of cell size.

## Discussion

Our results uncover previously unknown genome-wide patterns of transcriptional modulation from mitosis through late G1, observing transcriptional spiking at the mitosis-G1 transition for approximately half of all active genes and intergenic enhancers (Fig. 2). Prior to this work, an implicit expectation has been that early transcription at the mitosis-G1 transition resumes in a manner starting from initially low levels and then increase with varying kinetics to achieve maximal levels later in in interphase. However, only some genes in prior studies appear to display these characteristics (Blobel et al. 2009; Zhao et al. 2011; Kadauke et al. 2012; Fukuoka et al. 2012; Caravaca et al. 2013), while others exhibit non-monotonic patterns of transcription over time (Anup Dey et al. 2009; Muramoto et al. 2010; Fukuoka et al. 2012; Caravaca et al. 2013). Our current results demonstrate that the initial rounds of gene transcription upon reversal of mitotic silencing exhibit higher activities across the genome, compared to late G1, providing an overall context for interpreting results from prior studies. In particular, Muramoto et al. showed a spike in transcriptional activity immediately after mitosis of the act-5 gene by live-cell imaging in Dictyostelium. In light of our findings that the mitosis-G1 transcriptional spike is shared by at least two developmentally distinct murine cell lines (Fig. S10), these results together suggest that a spike in transcriptional activity after mitosis does not reflect a peculiarity of a specific gene or cell type, but is a general phenomenon of the genome that can be observed in evolutionarily distant cell types. The strength of our observation is supported by genome-wide coverage, purity of cells from the relevant cell cycle stages, extraction of prominent transcriptional patterns by unsupervised pattern discovery, and evidence that the mitosis-G1 transcriptional spike propagates to heterogeneity in single-cell mature mRNA levels.

What might be the mechanistic underpinning of the mitosis-G1 transcriptional spike? An important consideration is the effect of mitosis on the bulk distribution of transcription regulators. For clarity of discussion, we use Pol II as an example to illustrate a likely and potentially generalizable biophysical process. Mitotic displacement of Pol II would be expected to increases the unbound fraction of Pol II that would subsequently be available to initiate transcription upon reversal of mitotic inhibition. Since transcription initiation is restricted to promoter regions, this process would likely produce a transient increase in the ratio of effective enzyme concentration to the available DNA substrate. Upon reversal of mitotic silencing, such global shifts in factor distribution might predispose much of the genome to transcriptional spiking by mass action. Given that many general transcription factors have genomic occupancy distributions similar to Pol II during interphase and are likewise displaced from mitotic chromatin, the above scenario almost certainly applies to some factor that exists at a limiting concentration for transcriptional initiation. Such global changes in effective concentrations would be difficult to test for experimentally.

The degree of locus-specificity observed for the mitosis-G1 transcriptional spike requires additional explanations. Fig. 4 suggests that regions with high H3K27Ac might be preferred by Pol II re-initiation, explaining their proclivity to transcriptional spiking. Numerous studies proposed or assumed that chromatin-associated molecular entities marking individual loci during mitosis would influence the subsequent reading of the genome by the transcriptional machinery at the mitosis-G1 transition. Hence, these entities are metaphorically alluded to as mitotic “bookmarks.” How is H3K27Ac—now a candidate bookmark uncovered by genome-wide association—specified at individual loci during mitosis? Levels of histone acetylation in general are thought to result from the dynamic equilibrium of histone acetyltransferase versus histone deacetylase activities. Immunofluorescence microscopy previously showed that most acetylated histone H3 is globally reduced during mitosis (Kruhlak et al. 2001). This may reflect the outcome of bulk redistribution of both histone acetyltransferases—including p300 and CBP, known to be responsible for depositing H3K27Ac (Tie et al. 2009)—and histone deacetylases away from chromatin in mitosis (Kruhlak et al. 2001). Thus, our observation of significant retention of H3K27Ac by ChIP-seq at many loci during mitosis is likely an exception with respect to overall depletion of histone H3 acetylation, and indicates that some level of histone acetyltransferase activity must remain and exert locus-specific effects during mitosis. How the activities of these enzymes are specified at individual loci during mitosis remains unexplored.

What might be the biological consequence of the mitosis-G1 transcriptional spike? It is unclear whether this phenomenon has been programmed to serve a biological function. Recent single-molecule RNA FISH studies of candidate genes in mammalian cells (Padovan-Merhar et al. 2015; Skinner et al. 2016) and genome-wide studies in Saccharomyces cerevisiae (Voichek, Bar-Ziv, and Barkai 2016) have shown that total transcriptional output for individual genes before and after DNA replication in S-phase are equal. Our single-molecule RNA FISH measurements of per-copy gene transcription for the early G1 transcriptional spike genes, Gata2 and Kit, are consistent with these prior observations. Thus, on a per DNA copy level, transcriptional activity is two-fold higher in G1 than in G2, an observation first proposed by Padovan-Merhar et al. as promoting transcriptional homeostasis in the face of increased DNA copy upon replication. This doubled transcriptional activity per DNA copy overall in G1 necessarily includes contributions from the mitosis-G1 transcriptional spike. Thus, at least a portion of the transcriptional compensation in G1 arises from an unknown mechanism that exerts the most effect at the mitosis-G1 transition, rather than acting uniformly throughout G1. Such a model would not be mutually exclusive with, and could act in concert with, other potential mechanisms previously suggested to contribute to transcriptional gene dosage compensation, such as the dampening of transcriptional output upon nascent DNA synthesis in S-phase (Padovan-Merhar et al. 2015; Voichek, Bar-Ziv, and Barkai 2016). To what extent a dysregulation in gene dosage compensation at the transcriptional level might influence cellular function remains an intriguing open question.

Regardless of whether the mitosis-G1 transcriptional spike serves any particular biological function, our analysis of mature mRNA expression levels in transcriptionally “on” vs. “off” cells (Fig. 5) suggests that the transcriptional spike does not occur uniformly across a cell population. The differences in mature mRNA levels between the transcriptionally “on” and “off” cells (Fig. 5) appear modest (1.5 fold for Myc and 1.9 fold for Gata2) in the context of the population’s overall >100-fold range in mature mRNA levels. However, our approach of imaging fixed cells cannot directly evaluate a cumulative effect size that might be extracted from observing the trajectories of mRNA production in live individual cells over multiple cell divisions. To illustrate this possibility, suppose upon division of a single cell, the early G1 transcriptional spike stochastically occurs in one daughter cell but not the other, perhaps on average resulting in a 1.9-fold difference of mature mRNA levels in those two cells. Such a difference, while moderate to begin with, might predispose one cell for higher probability of subsequent higher expression levels. Such a scenario is consistent with, though not proven by, our indirect inferences from static images in Fig. 5, and might be particularly applicable if the gene product is involved in an autoregulatory positive feedback loop. Repeated sampling of the mitosis-G1 transition over multiple cell divisions might account for at least part of the eventual substantial divergence in gene expression state among all the progeny of the original founding cell. Consideration of the mitosis-G1 transition as a source of gene expression heterogeneity might pave the way for understanding why the probability of certain types of phenotypic transitions are modified by rapid proliferation (Smith et al. 2010), and passage through mitosis (Ganier et al. 2011; Halley-Stott et al. 2014; Egli, Birkhoff, and Eggan 2008) or early G1 phase (Singh et al. 2013). We envision that the mitosis-G1 transcriptional spike may, on average, promote gene expression homeostasis with respect to DNA dosage, yet its variable occurrence at the single-cell level may contribute to diversification of gene expression states.

## Experimental Procedures

### Cell culture, cell cycle synchronization, and cell sorting

G1E erythroblasts were previously derived through deletion of GATA1 in mouse embryonic stem cells, followed by in vitro differentiation (Weiss, Yu, and Orkin 1997). We cultured a sub-line of G1E cells, G1E-ER4, in which GATA1-ER was retrovirally transduced (referred to in main text as “G1E GATA1-ER”), as previously described (Weiss, Yu, and Orkin 1997). We retrovirally transduced G1E-ER4 cells with the YFP-MD construct (Kadauke et al. 2012) and sorted for a pool of YFP-positive cells. Except where indicated in the text as uninduced, we induced cells to mature with 100nM estradiol to activate GATA1-ER. During estradiol induction, we simultaneously treated cells with nocodazole (200ng/ml) for 7h-13h, washed once, and replated into fresh medium lacking nocodazole for varying times (40min-360min), ensuring all samples are exposed to estradiol for the same duration of 13h. We fixed cells with 1% formaldehyde, stained with DAPI, and sorted on a BD FACSAria based on YFP-MD and DAPI signal. Sorting of MPM2-positive prometaphase populations and MPM2-negative interphase populations for H3K27Ac ChIP-seq was carried out as described in (Campbell, Hsiung, and Blobel 2014).

F9 cells (ALONSO’ et al. 1991) were cultured in plates pre-coated with 0.1% gellatin and grown in DMEM + 10% FBS. For mitotic arrest-release, cells were treated with nocodazole (200ng/ml) for 4h and a “shake-off” (gentle rinsing with media) was performed to isolate mitotic cells, followed by replating into fresh media for varying durations of the release time course.

### Chromatin immunoprecipitation-sequencing

We performed ChIP-seq of total 3 biological replicates using N-20 antibody (Santa Cruz, cat# sc899) for the 0min, 60min, 90min, 180min, and 360min time points; two biological replicates for 240min time point; and one replicate for 40min time point. Two replicates of input DNA at the corresponding time points were also sequenced. For ChIP-qPCR of initiating form of Pol II, we used 8WG16 antibody (Covance, cat# MMS-126R). H3K27Ac antibody from ActiveMotif (cat# 39685) was used for H3K27Ac ChIP-seq. To summarize briefly, cells fixed with 1% formaldehyde and subjected to lysis in detergents, sonication, and immunoprecipitation of chromatin. Following library construction through blunt end repair and adaptor ligation using Illumina’s TruSeq ChIP Sample Preparation Kit (Illumina cat# IP-202-1012), size selection with SPRIselect Beads (Beckman Coulter cat# B23318), and PCR amplification, libraries were multiplexed and sequenced on the Illumina HiSeq 2000. The mean fragment size is approximately 330bp. See Supplemental Experimental Procedures for details.

### Bioinformatic analysis of ChIP-seq data

To summarize briefly, reads were mapped to mouse mm9 genome using Bowtie (Langmead et al. 2009). Mapped reads were passed to MACS (Zhang et al. 2008) with a matched control (input) dataset for peak calling and for producing bigwig files with reads shifted to account for fragment size. If the 5’ or 3’ 2.5kb region of a gene overlapped at least one Pol II peak called by MACS in at least one sample (arrest-release and asynchronous samples with estradiol induction), then we deemed the gene active, arriving at 4309 non-overlapping active genes defined by the single largest annotated transcript of each gene. We defined 809 intergenic enhancers as those DNase hotspot regions previously described to overlap H3K4me1 in the relative absence of H3K4me3 (Hsiung et al. 2014), overlap Pol II MACS peak, and must not overlap the 5’-3kb and 3’ +10kb regions of annotated genes. “Pol II binding” or “read density” used in the context of Pol II ChIP-seq in the text refer to reads per million kilobases per million mapped reads (RPKM) from regions of interest. We performed principal component analysis on RPKMs normalized by the sum of RPKM for each gene across G1 time points (60min, 90min, 180min, 240min for replicates 2 and 3, and 360min) using the R package prcomp and custom scripts. Details are provided in Supplemental Experimental Procedures, and scripts provided as described in the Data Access section.

### RT-qPCR of primary transcripts

We isolated RNA using TRIzol (Life Technologies) or TRIzol LS (Life Technologies). Reverse transcriptase reaction was performed with iScript (Bio-Rad). qPCR was performed with Power SYBR Green (Invitrogen). All primer sequences are provided in Supplemental File. For primary transcript measurements, primers flank intron-exon junctions. Primary transcript quantifications are normalized to Gapdh mature mRNA. Results are similar when normalized to Hprt mature mRNA (not shown).

### Capture-C

After cell fixation with 1% formaldehyde for 10min and cell sorting as described above, Capture-C was performed with a double-capture procedure (Davies et al. 2016). Chromatin was digested using DpnII. We used biotin labelled DNA oligos to pull down the target restriction fragments. Capture-C libraries were sequenced on Nextseq500 with 2x75 bp paired end sequencing. The raw reads were processed using published scripts (https://github.com/telenius/captureC/releases). We wrote custom scripts to normalize data by the total number of reads representing fragments ligated to the anchor region in the library. Enhancer regions used for quantitation were defined by assessment of Capture-C signal by eye, together with consideration of prior literature and DNase sensitivity signals.

### Single-molecule RNA FISH

We performed single-molecule RNA FISH as described previously (Femino 1998; Raj et al. 2008; Raj and Tyagi 2010; Levesque and Raj 2013). In brief, we fixed cells in 1.85% formaldehyde for 10min at room temperature, and stored them in 70% ethanol at 4deg. C until further processing. FISH probes consist of oligonucleotides conjugated to fluorescent dyes as follows: Myc exons to Cy5, Gata2 exons to Cy3, Myc introns to Alexa594, and Gata2 introns to Alexa594. Oligonucleotide sequences are provided in Supplemental File. Imaging was performed on a Nikon Ti-E inverted fluorescence microscope using a 100x Plan-Apo objective (numerical aperture of 1.43), a cooled CCD camera (Pixis 1024B from Princeton Instruments), and filter sets SP102v1 (Chroma), SP104v2 (Chroma), and 31000v2 (Chroma) for Cy3, Cy5 and DAPI, respectively. Custom filter (Omega) was used for Alexa594. We took optical z-sections (typically 45) at intervals of 0.35 microns, spanning the vertical extent of cells, with 1s exposure time for Cy3, Cy5, and Alexa594, and 100ms for DAPI.

### Image analysis

We manually segmented boundaries of cells from brightfield images and localized RNA spots using custom software written in MATLAB (Raj and Tyagi 2010). The area within segmentation borders is used for cell area. For Fig. 6, we adjusted for minor systematic variations in the distributions of cell area found across imaging sessions by adding a constant to the cell area, such that the median across all biological replicates are equal. Mature mRNA concentrations per cell are quantified by the spot counts in the exon channel, divided by the cell area. Primary transcripts are identified by intron and exon co-localization as detailed in Supplemental Experimental Procedures. For Fig. 5, actively transcribing cells are defined as those that have at least one intron-exon colocalized spot.

### Data Access

All raw and processed sequencing data will be deposited at GEOXXXXXX. In addition, scripts that reproduce the majority of figures starting from processed data in tabular form (including RPKMs from ChIP-seq, spot counts and intensities from single-molecule RNA FISH, read counts from Capture-C, and CT values from RT-qPCR) are provided and maintained in an online repository

(https://chsiung@bitbucket.org/arjunrajlaboratory/hsiung_mitosisreactivation), as well as in a Supplemental File. Pol II ChIP-seq tracks can be viewed at genome browser hosted by Pennsylvania State University (http://main.genome-browser.bx.psu.edu/cgibin/hgTrackUi?hgsid=206746_6ux0ZS2v9DeKCbVJzcfJ2tRchNFS&c=chr5&g=meryYfpmd

### Author Contributions

C.C.H., A.R., and G.A.B. planned the study. C.C.H., C.B., P.H., A.J.S., C.A.K., C.F., K.S.J., and L.S. performed experiments. C.C.H, C.B., P.G., P.E., P.H. and B.G. performed computational data processing and analyses. C.C.H., R.C.H., A.R., G.A.B. interpreted the results. C.C.H., A.R., and G.A.B. wrote the manuscript with input from all authors.

**Fig. 7:**
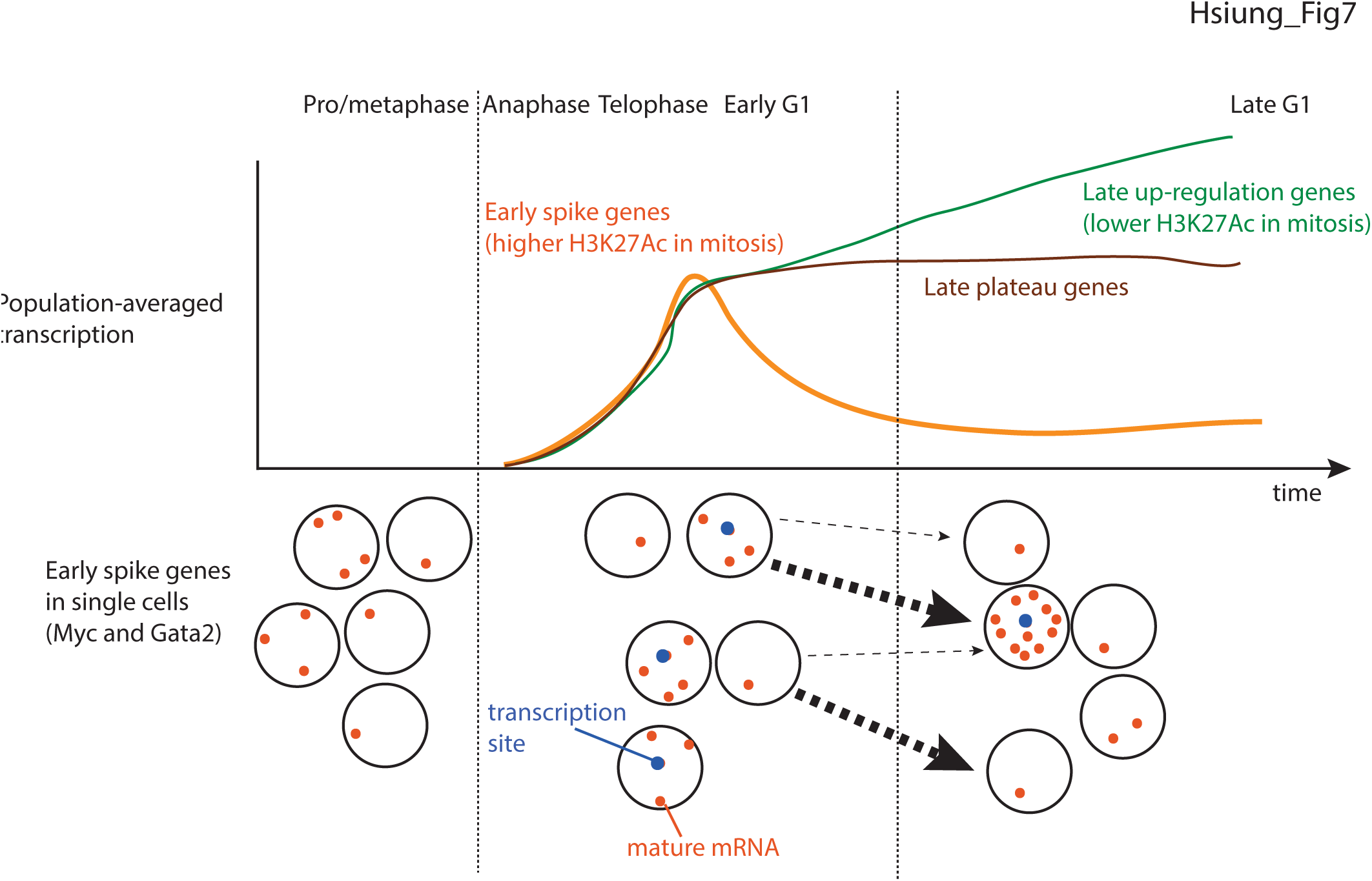
Model of transcriptional patterns in mitosis and G1 phase. Summary of genome-wide transcriptional patterns during progression from mitosis through G1 phase on the population-averaged level, and on the single-cell level for early spike genes. In the single-cell illustration, arrows represent likely single-cell transitions over time, with the sizes of the arrows qualitatively representing the relative probabilities of those transitions.

## Acknowledgement

We thank Hua-Ying Fan and Robert Lake for generous gift of F9 cells. We thank Sarah Hsu, Katherine Palozola, Sheila Teves and Kenneth Zaret for critical reading of the manuscript. We also thank Gautham Nair, Marshall Levesque, Jennifer Phillips-Cremins, Michael Lampson, Hua-Ying Fan and Stephen A. Liebhaber for helpful discussions.

## Supplemental Results

**Figure S1.**
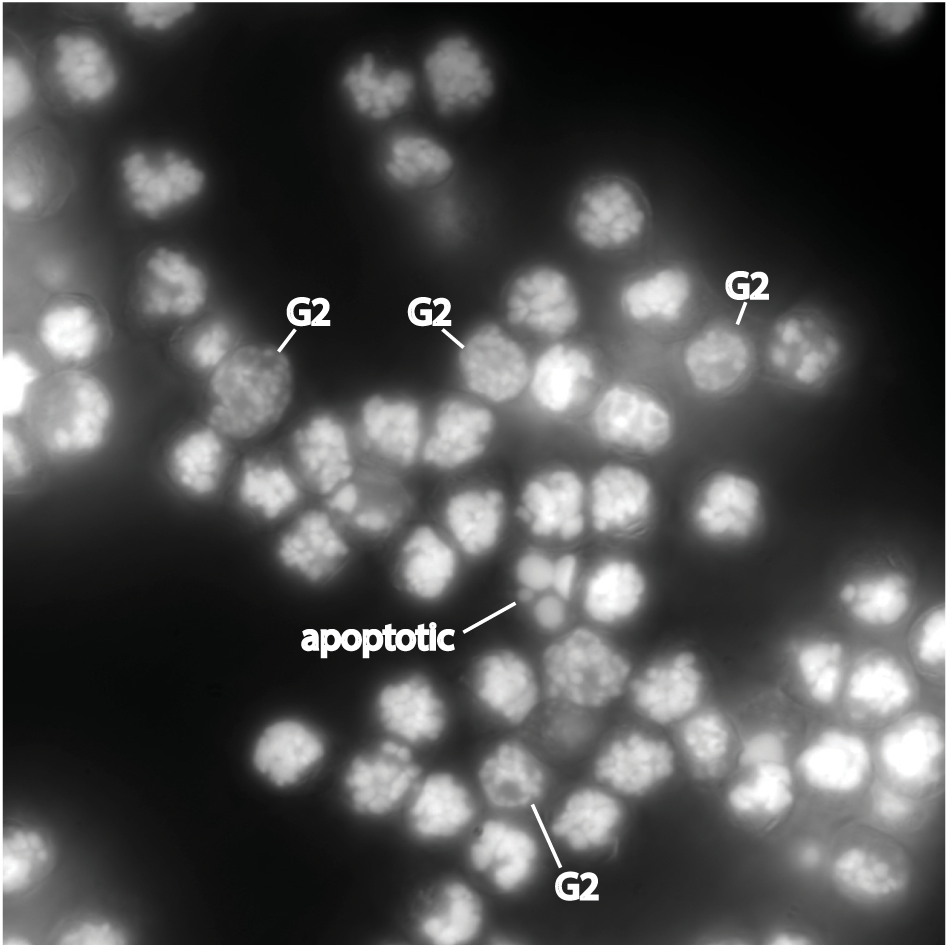
Related to Fig. 1A: Estimate of interphase contamination in 0min sample. Cells were arrested with nocodazole and sorted for 4N, YFP-MD-high as described in Fig. 1A for the 0min time point. Shown is a representative field of cells in the DAPI channel by wide-field microscopy (single optical plane is shown). There is approximately 10% G2 contamination (27 out of 271 cells), as judged by microscopic appearance of DAPI staining, and minimal apoptotic cells.

**Figure S2.**
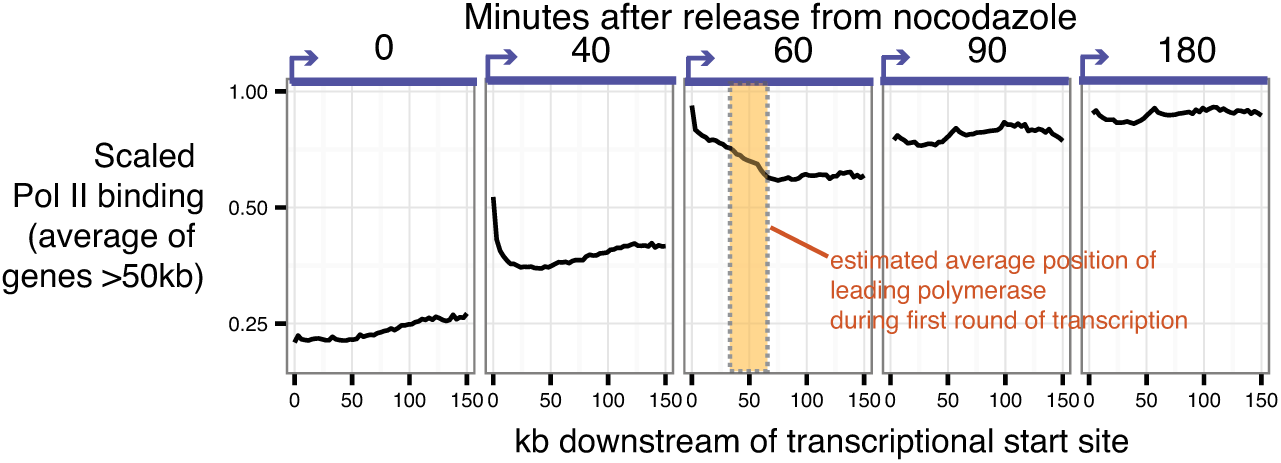
Related to Fig. 1B: Genome-wide average position of Pol II leading edge. Average Pol II ChIP-seq read densities (scaled to the read density in the asynchronous sample of each gene) obtained from 3kb bins across all genes >50kb is plotted in the sense direction along genomic coordinates relative to the transcriptional start site. The estimated averaged position of the Pol II leading edge at the 1h time point, determined manually by eye, is indicated by the shaded region.

**Figure S3.**
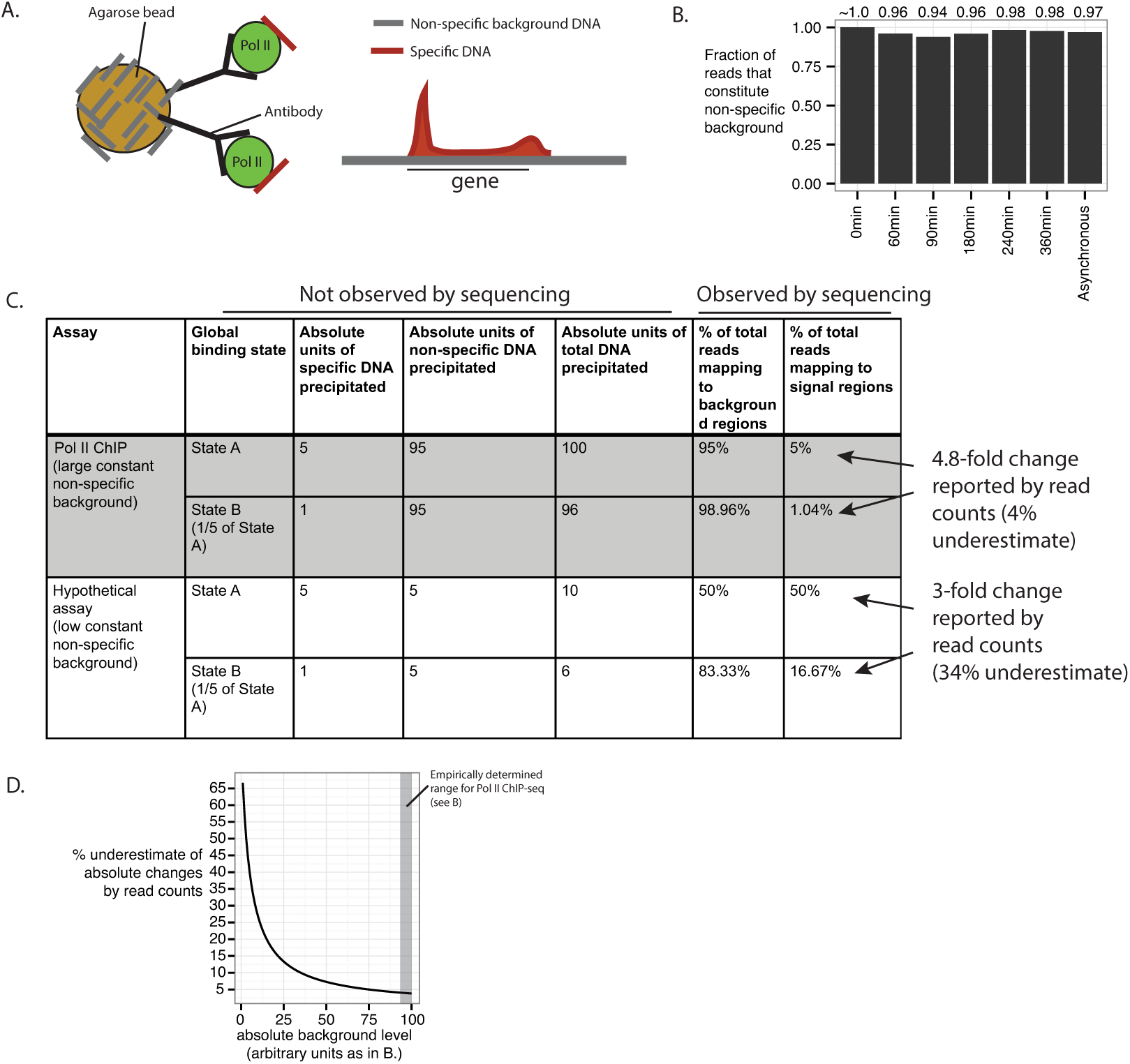
Related to Fig. 2: Pol II ChIP-seq signal in genes is internally normalized to reads mapping to non-specific background, enabling interpretations of absolute binding. **A)** Model of the physical origins of Pol II ChIP-seq reads that map to genic (regions defined by Refseq annotations), versus intergenic regions (outside of Refseq annotations). Reads mapping to intergenic background regions likely arise from DNA bound non-specifically to surfaces of reagents used to pull-down the antibody. We assume that the amount of non-specifically bound DNA is constant across samples processed identically. **B)** The fraction of total mapped reads that constitute intergenic non-specific background empirically determined for each Pol II ChIP-seq library is >94%. Note that the slight dip at 60min and 90min time points is consistent with the overall spike in Pol II binding within genes described in main text. **C)** We illustrate the effect of the level of non-specific background on the ability of a sequencing-based assay to estimate absolute changes in signal. We consider a biological phenomenon that causes a global 5-fold change in Pol II binding from State A to State B, and assess the performance of our Pol II ChIP assay in reporting this change, compared to a hypothetical assay in which absolute level of non-specific background is 19 fold less. This analysis shows that our Pol II ChIP-seq would underestimate the true fold change by only 4%, whereas the hypothetical low-background assay would underestimate the fold change by 34%. **D)** For the same scenario as in C, we illustrate the relationship between error in estimating absolute changes by library size-normalized read counts with constant absolute background inherent to the assay.

**Figure S4.**
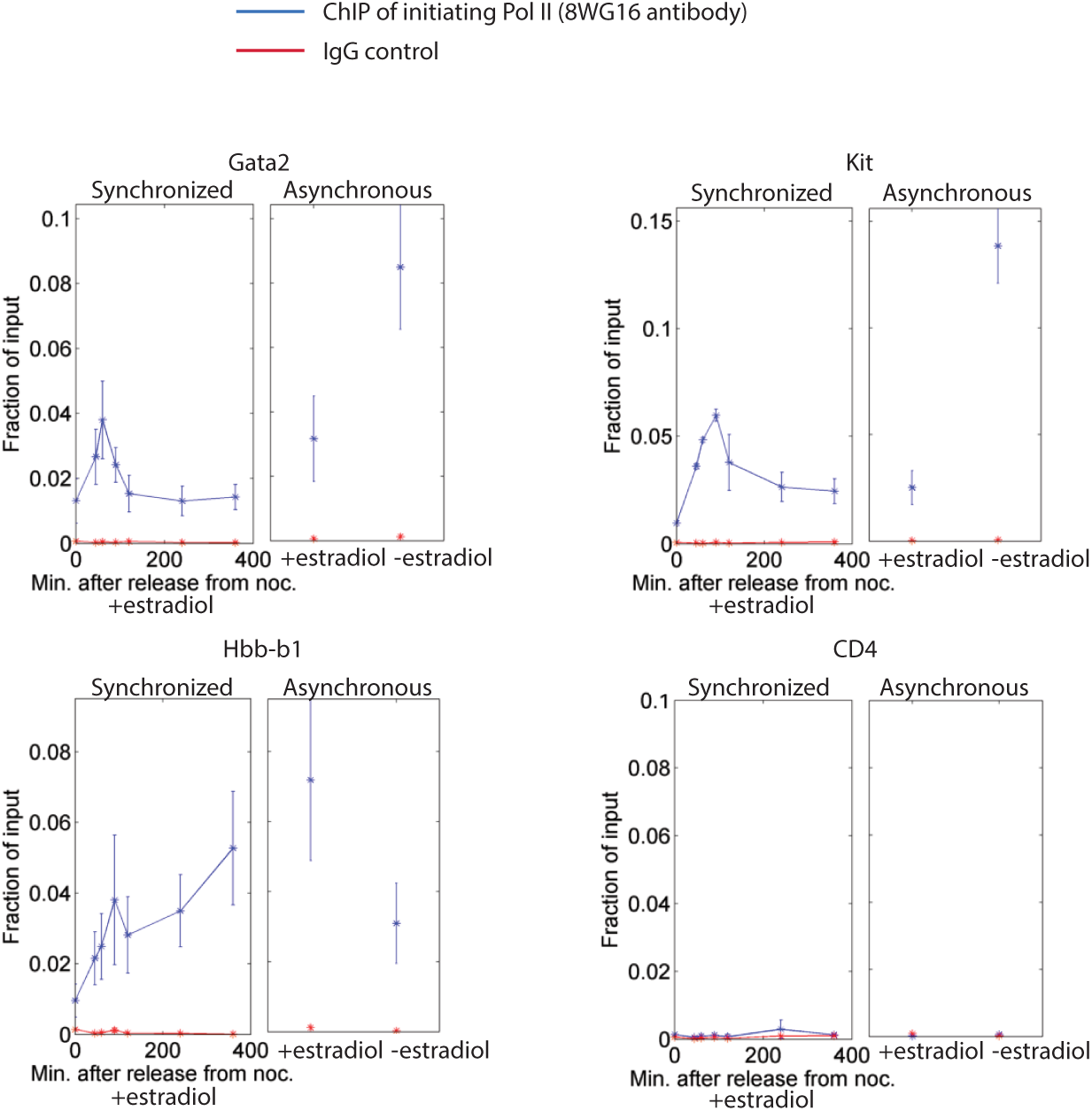
Related to Fig. 2B: ChIP-qPCR of initiating Pol II confirms ChIP-seq patterns at individual loci. We performed ChIP using an antibody (8WG16) specific for the initiating form of Pol II in estradiol-induced G1E GATA1-ER cells, followed by qPCR of amplicons proximal to the transcriptional start site in a nocodazole arrest-release time course without additional FACS purification. Also shown are ChIP performed in asynchronous controls with and without estradiol treatment. Gata2 and Kit are known to be down-regulated by estradiol induction, Hbb-b1 is known to be up-regulated by estradiol induction, and CD4 is a silent gene that serves as a negative control. Error bars denote SEM (n=3-4)

**Figure S5.**
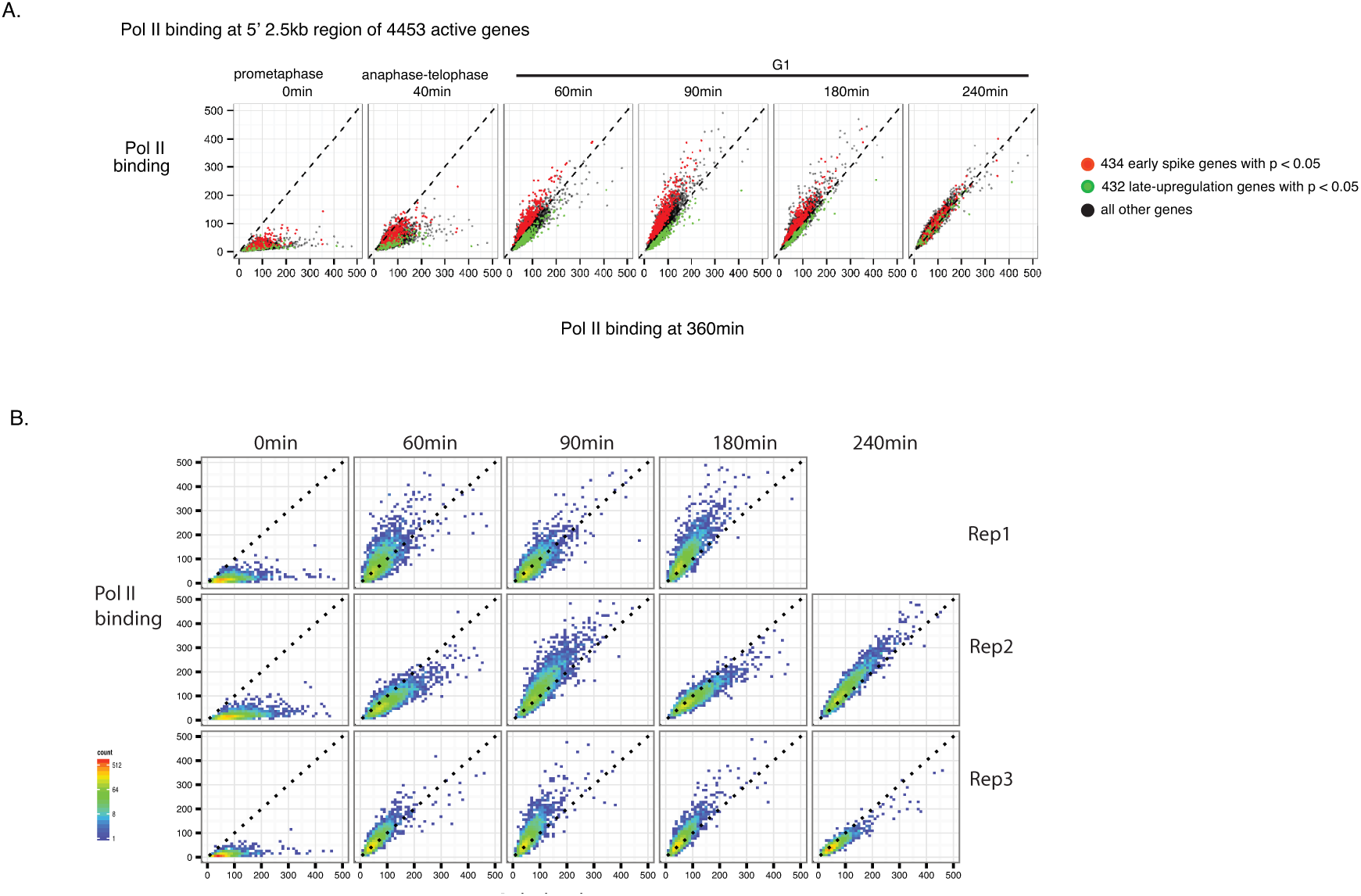
Related to Fig. 2: Statistical significance of degree of match to the first principal component for individual genes and genome-wide Pol II binding patterns for each biological replicate. **A)** A bootstrapping approach was used to assess statistical significance of the degree of match to the first principal component for each gene. An empirical null distribution was obtained by scrambling the time points for each gene 10000 times without replacement and projecting the scrambled data onto the the first principal component eigenvector. Genes with p < 0.05 based on a one-sided test are highlighted in the graphs showing Pol II binding at each time point plotted against that at the 360min time point. B) Pol II binding at the 5’ 2.5kb region of 4309 genes active in at least one time point were plotted for each time point against the 360min time point for each of the biological replicates.

**Figure S6.**
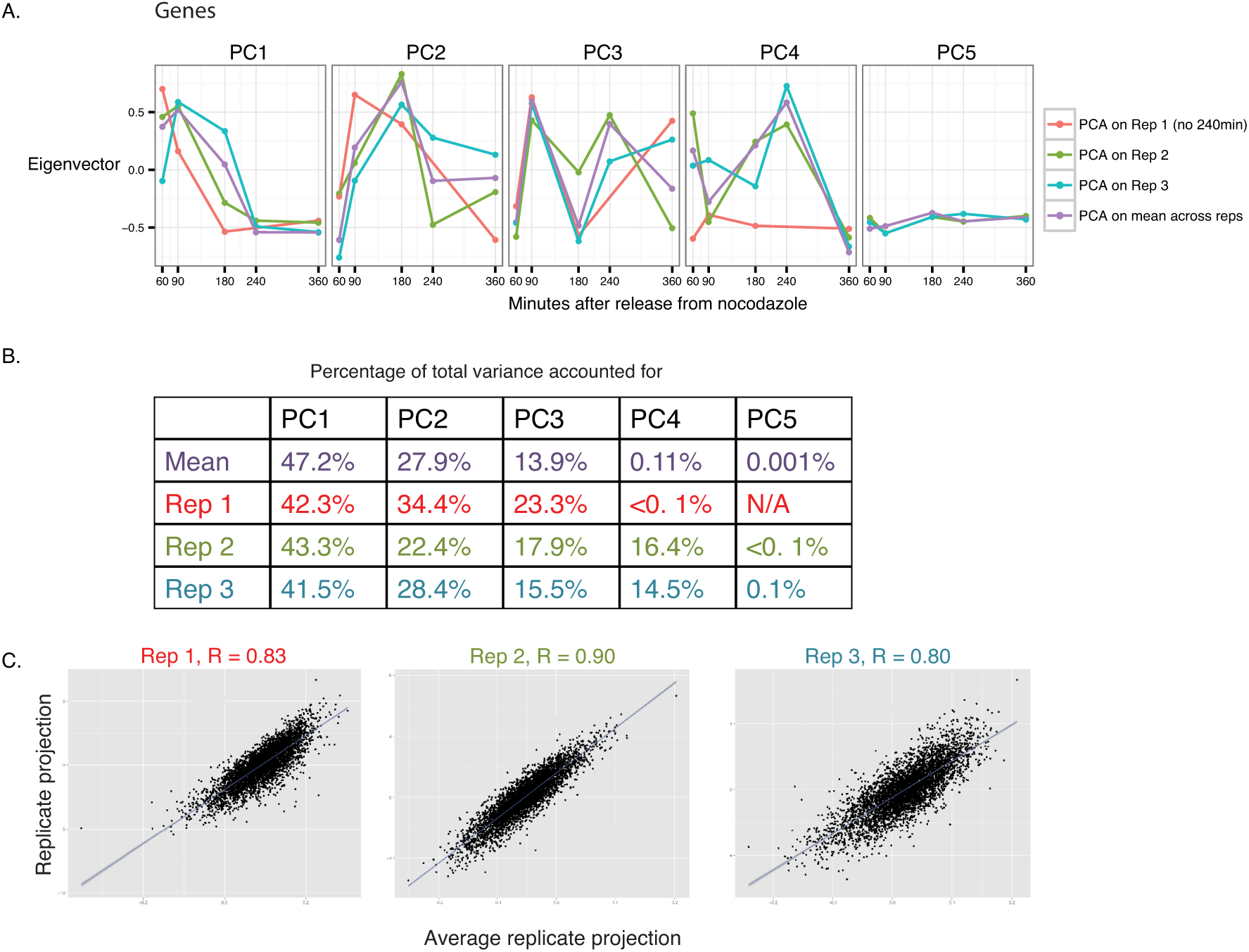
Related to Fig. 2B: Principal component analysis for Pol II binding at genes. **A)** We performed principal component analysis on each of the three biological replicates using the 5’ 2.5kb region of 4309 genes deemed active in at least one of the time points. Only G1 time points (60min, 90min, 180min, 240min, 360min) were used for principal component analysis. Shown are the principal components found for each replicate. Principal components found by using the mean Pol II binding across all replicates are also shown (same first principal component as shown in Fig. 2B). Replicate 1 does not have the 240min time point, so has only 4 principal components. **B)** The percentages of total variance accounted for by each of the replicate-derived and mean-derived principal components are shown. **C)** We evaluated replicate concordance by plotting the projection of the RPKM for each replicate onto the first principal component derived from the mean RPKM across all replicates on the y-axis, versus the projection of the mean RPKM across all replicates onto the first principal component derived from the mean RPKM across all replicates on the x-axis. Pearson correlation coefficients are shown.

**Figure S7.**
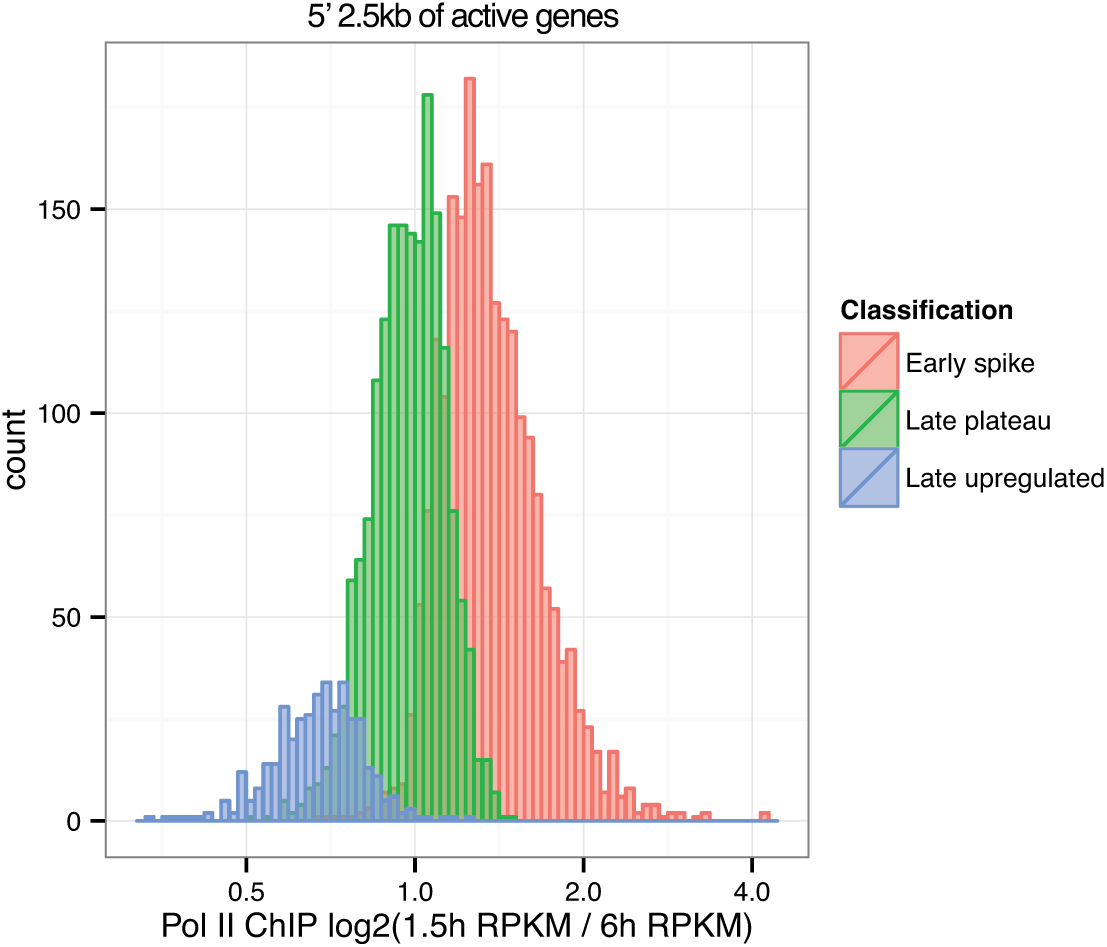
Related to Fig. 2B: Magnitudes of transcriptional changes in G1. Pol II binding is quantified as ratio of RPKM from 1.5h time point to the 6h time point, and shown as histograms separately for the early spike, late plateau, and late upregulated classes defined in Fig. 2B.

**Figure S8.**
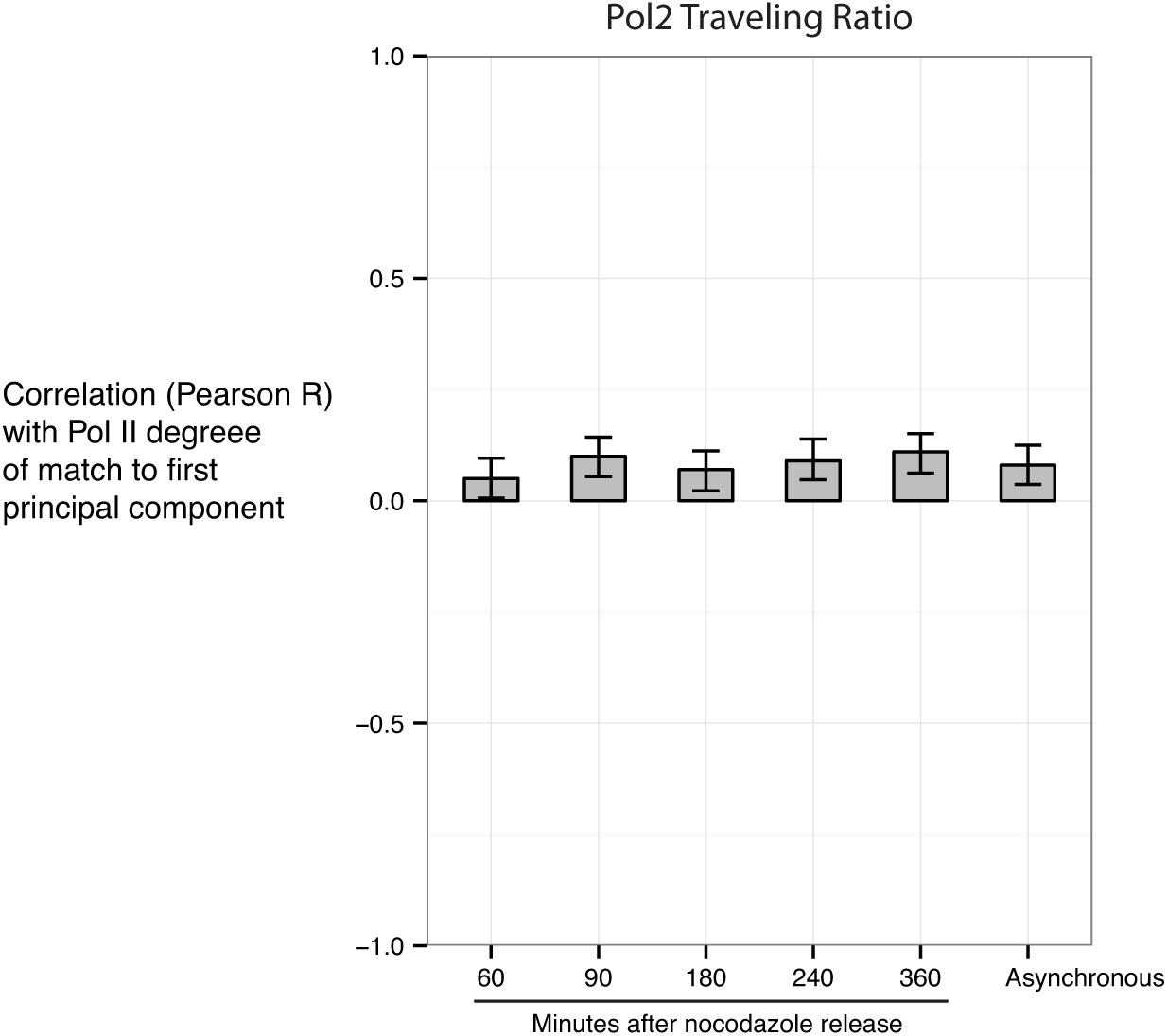
Related to Fig. 2: Mitosis-G1 transcriptional spike is not associated with changes in Pol II promoter escape. Pol II degree of match with (projection onto) the first principal component is plotted against a traveling ratio of Pol II (a measure of the rate of Pol II promoter escape) at each time point. We modified the definition of the traveling ratio (Rahl et al., 2010) to be the Pol II ChIP-seq read density in the −30bp to +300bp region relative to the TSS, divided by the read density in the +1.3kb to +5kb region. This definition avoids confounding from incomplete elongation in the early time points after mitosis. The analysis was performed only on genes >5kb.

**Figure S9.**
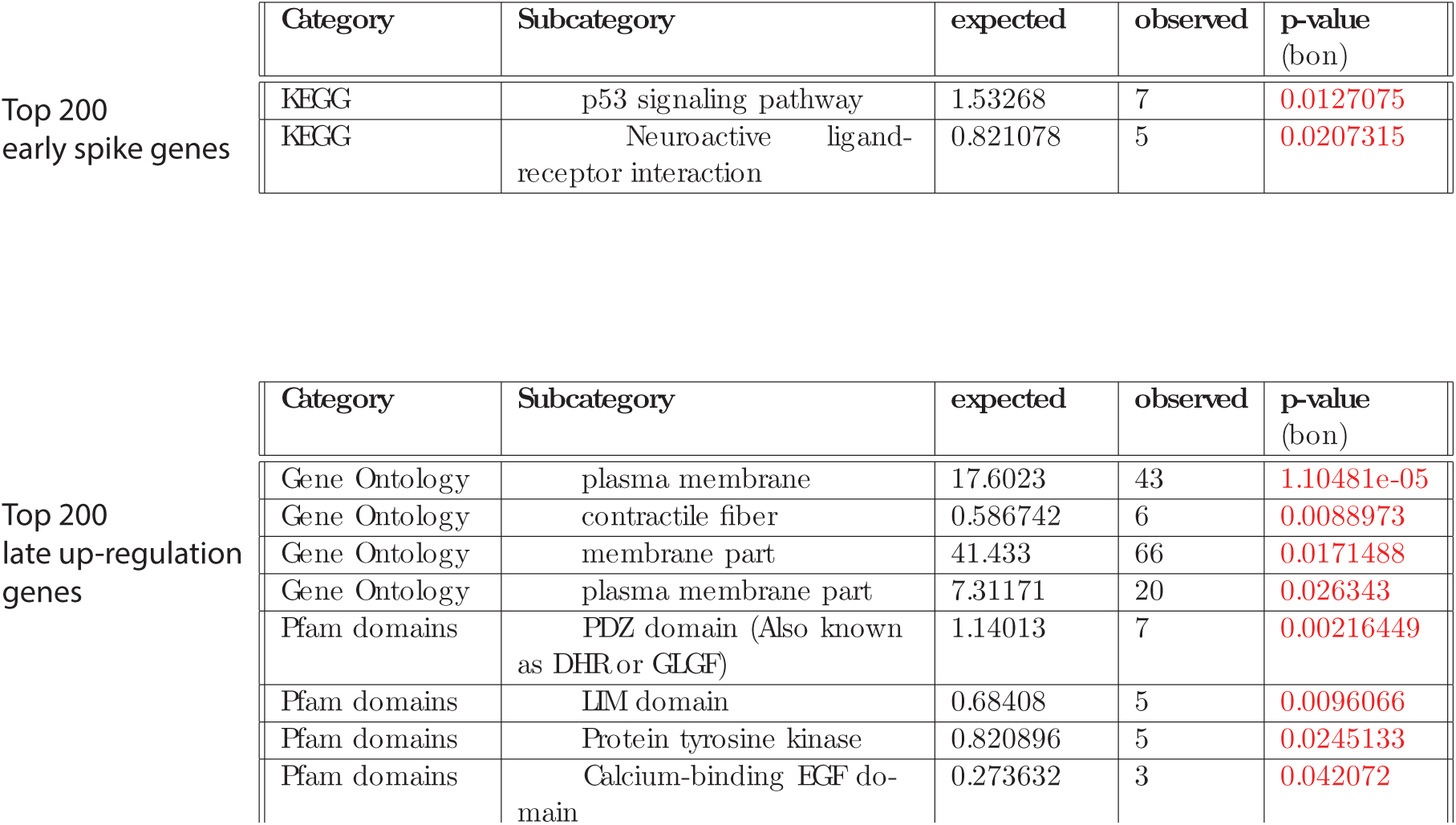
Related to Fig. 2B: Analysis of gene sets enriched among early spike vs. late up-regulalion genes. Using GeneTrail (Backes el al., 2007; Keller el al., 2008), we tested for gene seis thai are overrepresented among the top 200 early spike genes or the top 200 late up-regulation genes, using the 4309 active genes as background. GeneTrail includes tests for the enrichments of KEGG pathways. Gene Ontology and Pfam domains. Shown are the overrepresented gene sets that are statistically significant (Bonferroni-corrected p = 0.05).

**Figure S10.**
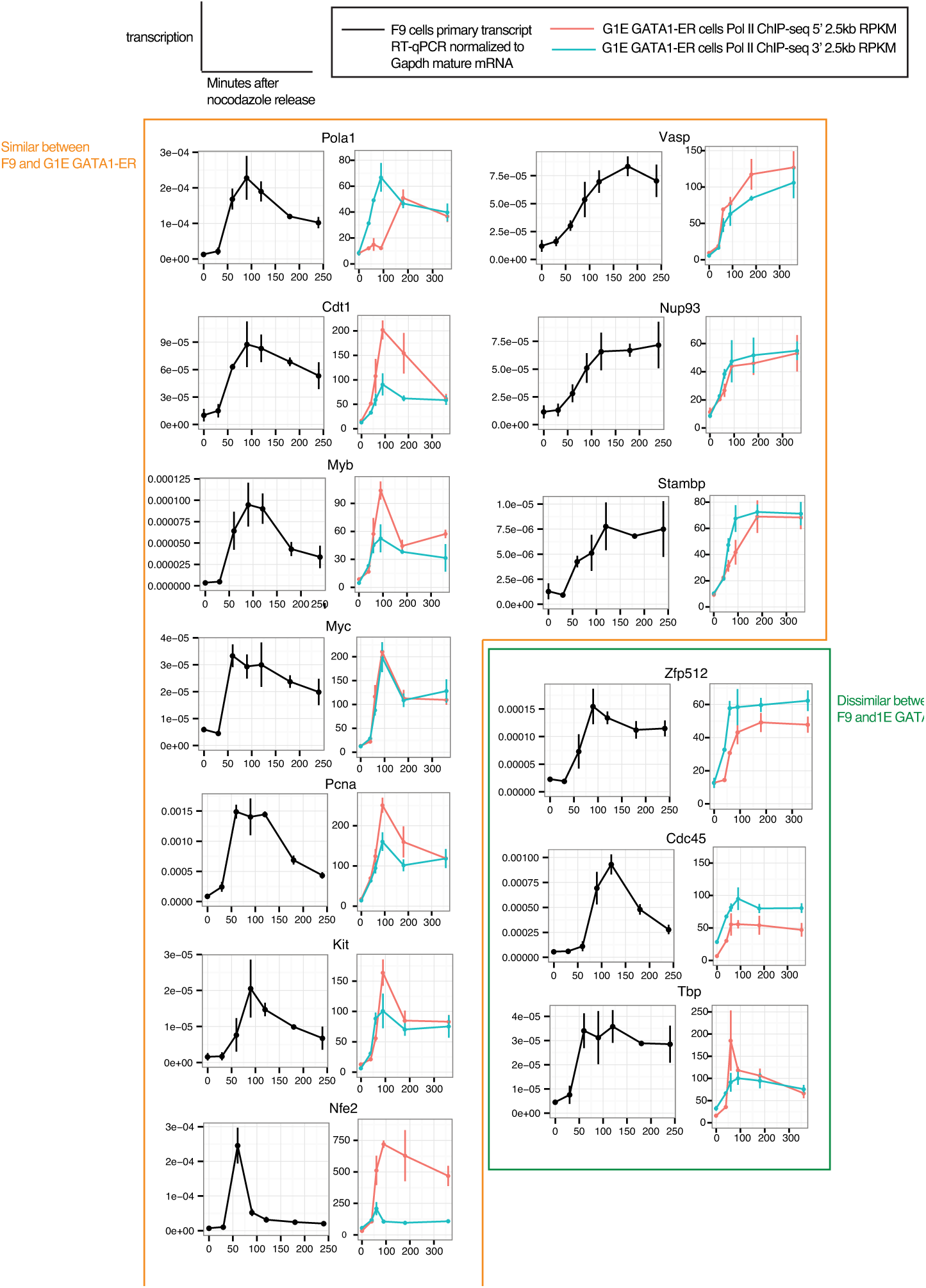
Comparison of G1 transcriptional patterns between developmentally distinct murine cell types. Shown are nocodazole arrest-release transcriptional profiles for an embryonic carcinoma cell line (F9) and erythroid cell line (G1E GATA1-ER). Genes expressed in both cell lines were examined by primary transcript RT-qPCR using primers flanking intron-exon junctions for F9 cells, and Pol II ChIP-seq for G1E GATA1-ER as described in the main text. Several of the G1E GATA1-ER Pol II ChIP-seq plots are duplicated from Fig. 1 and Fig. 3 for ease of comparison. Genes showing similar or dissimilar G1 transcriptional profiles are highlighted. E1r2ror bars denote SEM (n = 4 for F9; n = 3 for G1E GATA1-ER).

**Figure S11.**
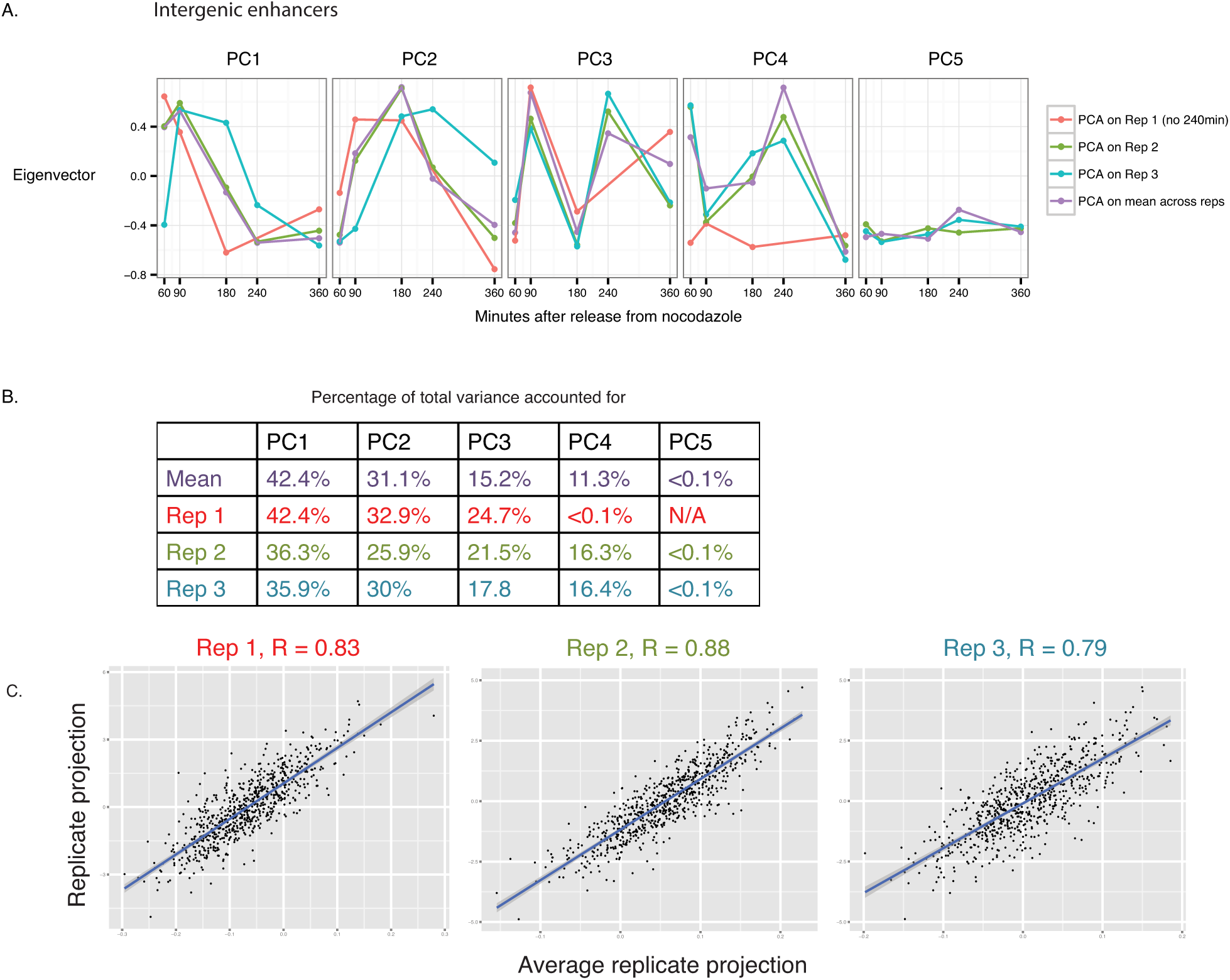
Fig. 3A: Principal component analysis for Pol II binding at intergenic enhancers. Principal components analysis was performed in for 809 intergenic enhancers in the same manner as described for genes in Fig. S6, and the information shown in A-C are exactly analogous to that in Fig. S6.

**Figure S12.**
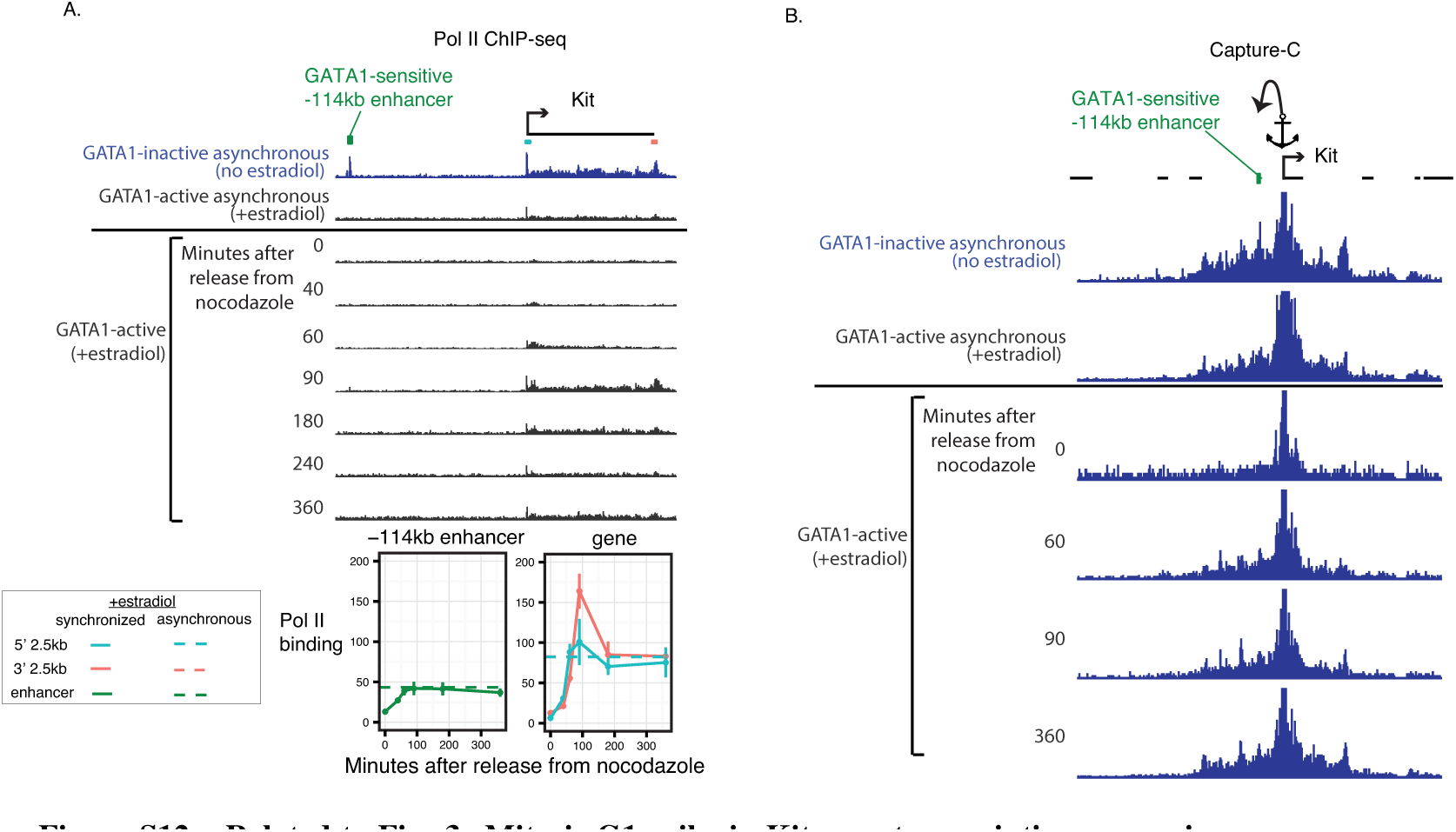
Related to Fig. 3: Mitosis-G1 spike in Kit gene transcription occurs in the absence of spike in Pol II binding or enhancer-promoter looping. **A)** Browser track views of Pol II binding at the Kit locus, including the enhancer located at −114kb upstream of the gene. Tracks are shown for GATA1-inactive condition (no estradiol treatment), in which there is high Pol II binding to the gene and enhancer, and for GATA1-active conditions (13h treatment with estradiol), in which there is basal level of Pol II binding at the gene and very little at the enhancer. A time course of release from nocodazole arrest is shown in the GATA1-active condition. Quantification of Pol II binding at the −114kb enhancer and the 5’ 2.5kb and 3’ 2.5kb regions of the gene are shown below. **B)** Browser track views of Capture-C signal performed with anchor at the Kit promoter, under conditions as described in A). Quantification of Capture-C enhancer-promoter contacts are shown in Fig. 3D.

**Figure S13.**
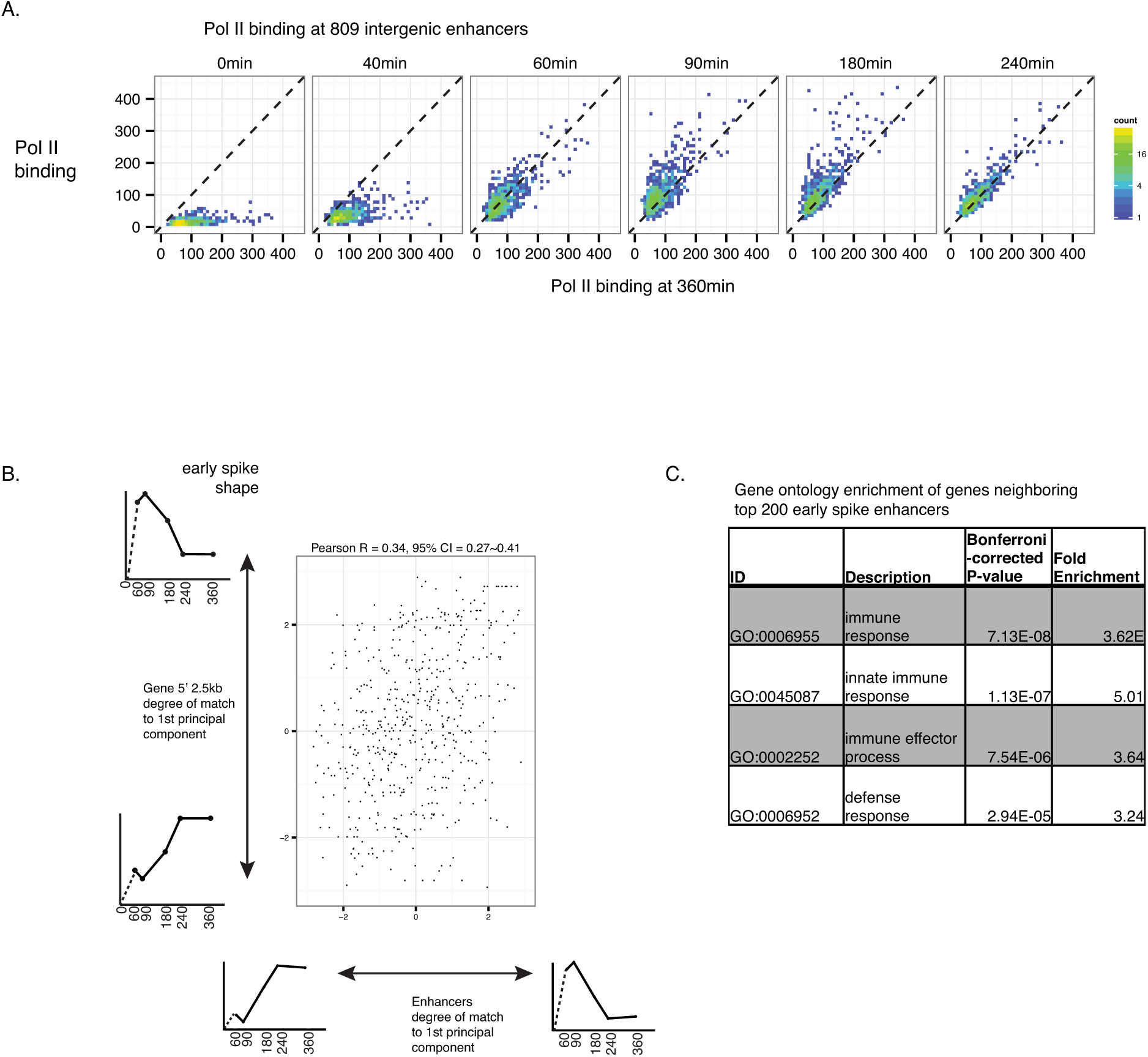
Related to Fig. 3: Global patterns of Pol II binding at intergenic enhancers and their nearest gene. **A)** Pol II binding at 809 intergenic enhancers at each time point is plotted against binding at the 360min time point. **B)** Each enhancer is assigned to the nearest gene. The degree of match to (projection onto) the 1st principal component for enhancers (as described in Fig. 3A) is plotted against the degree of match to (projection onto) the 1st principal component of the nearest gene (as described in Fig 2B). The Pearson correlation coefficient is shown. C) We used GREAT 3.0.0 McLean et al. (2010) to assess Gene Ontology enrichments among neighboring genes of the top 200 early spike intergenic enhancers, using the set of 809 intergenic enhancers as background. Shown are enriched ontologies with at least 6 genes associated with the target regions and have p < 0.05 after Bonferroni correction.

**Figure S14.**
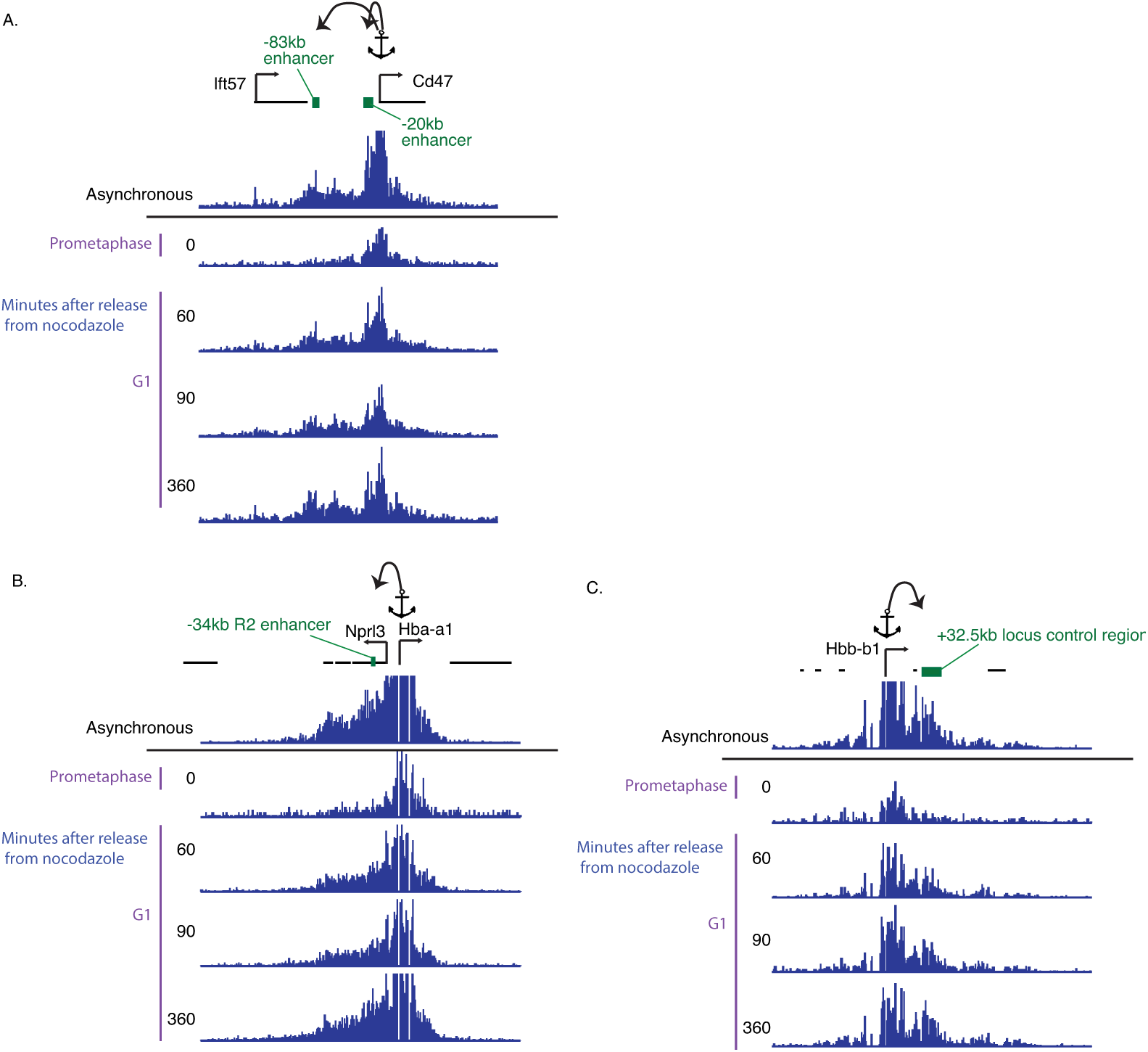
Related to Fig. 3: Browser tracks for enhancer-promoter contacts at the mitosis-G1 transition measured by Capture-C. Capture-C was performed using anchors targeting the A) Cd47 promoter, B) Hba-a1 promoter, and C) Hbb-b1 promoter, as described in Fig. 3C. The corresponding quantification of read densities at individual enhancers are shown in Fig. 3C.

**Figure S15.**
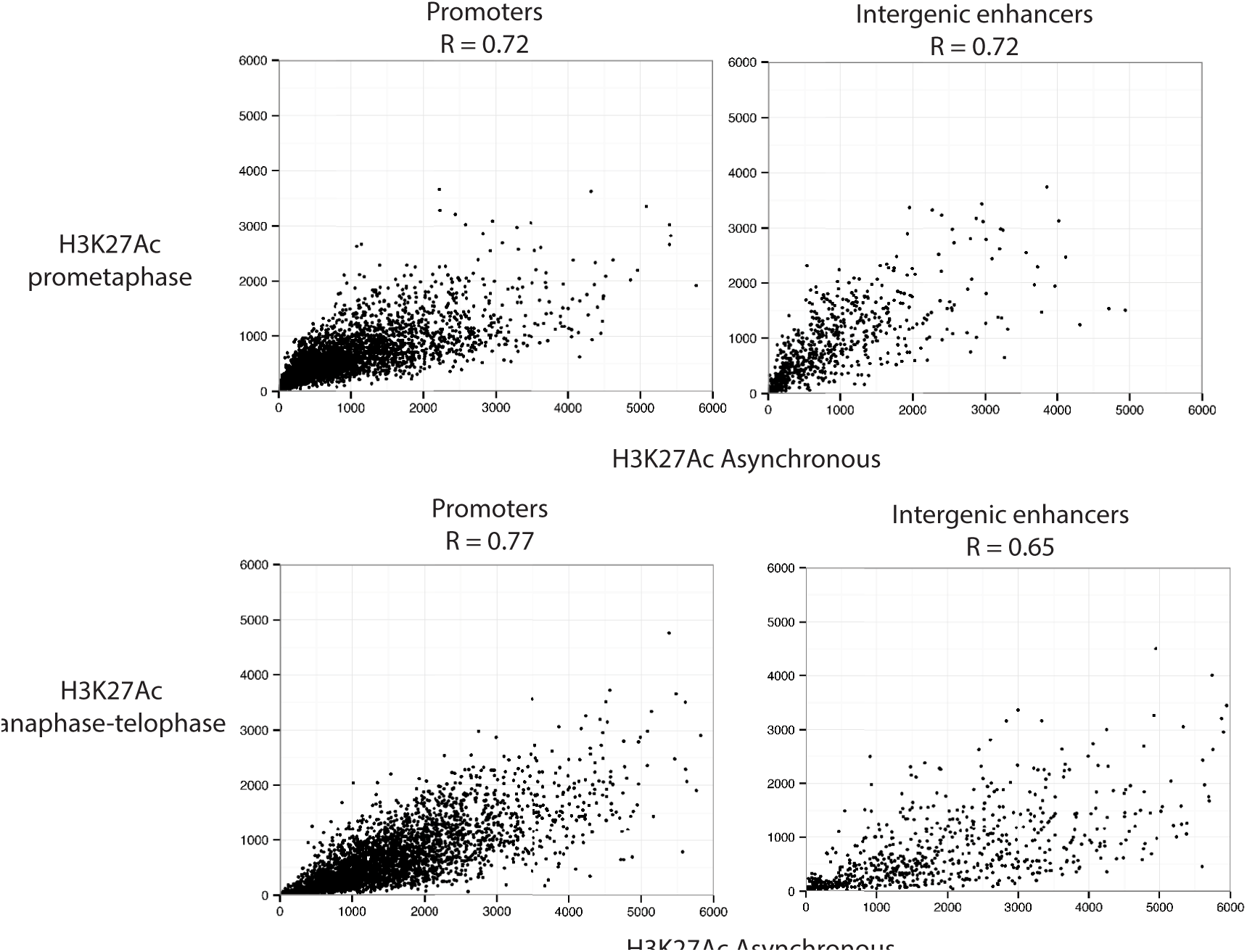
Related to Fig. 4: Locus-specific differences in H3K27Ac in mitosis and interphase. Top row: Scatter plots of H3K27Ac ChIP-seq for cells arrested by nocodazole in prometaphase (FACS purified for MPM2-positivity) vs. asynchronous (FACS purified for MPM2-negativitiy). Bottom row: Scatter plots of H3K27Ac ChIP-seq for cells in the anaphase-telophase compartment (40min after release from nocodazole arrest, sorted for YFP-MD low and 4N DNA content by DAPI) vs. matched asynchronous sample. Each data point corresponds to a DNase hotspot annotated as promoters or intergenic enhancers. Values shown are library size-normalized read densities (RPKM). Pearson correlation coefficients are shown at the top of each graph.

**Figure S16.**
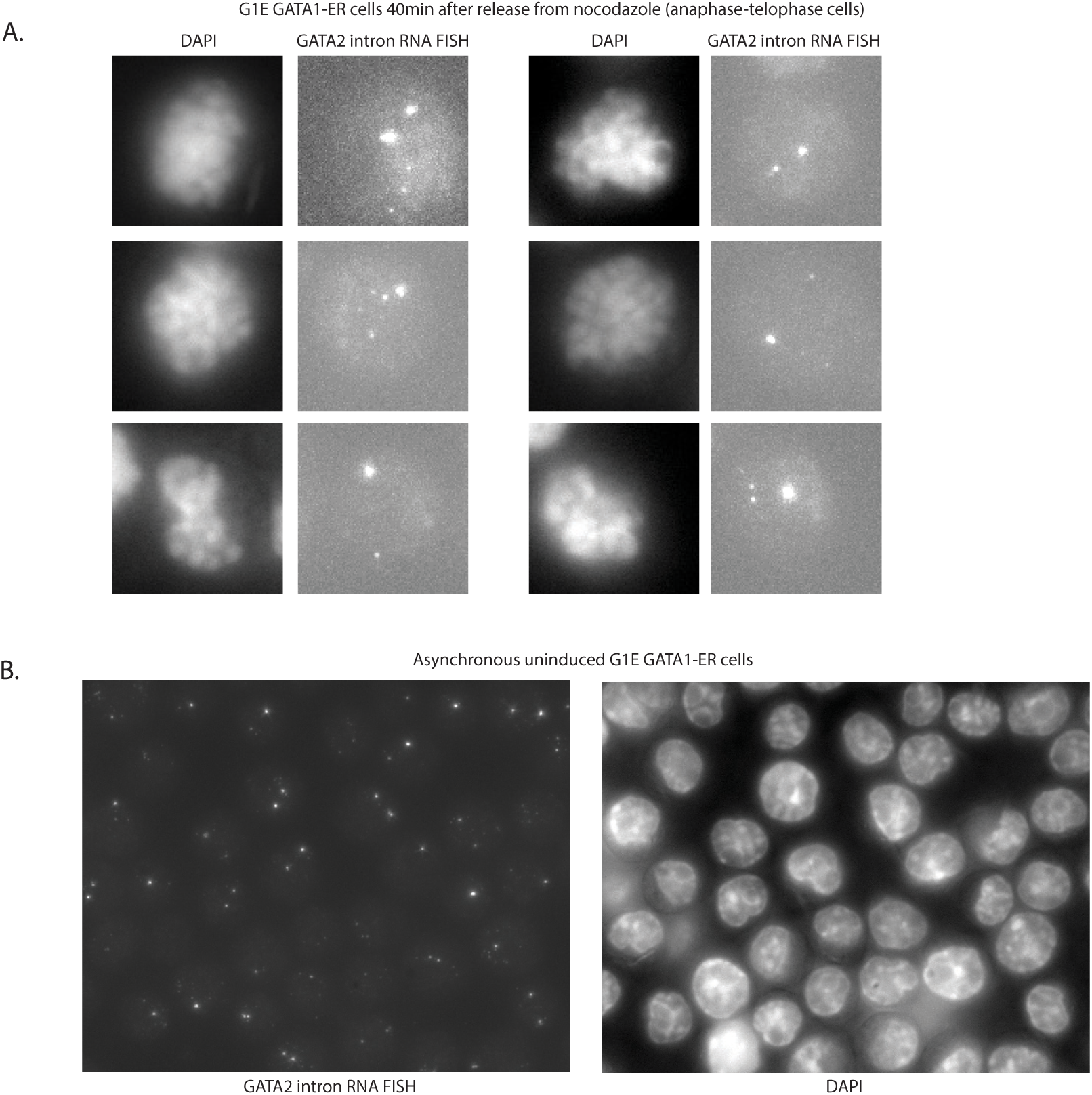
A Additional images of anaphase-telophase G1E GATA1-ER cells (estradiol induced, 40min after release from nocodazole) with grossly condensed chromosomes and active transcription detected by single-molecule RNA FISH for GATA2 introns. B) Images of an asynchronous population of uninduced G1E GATA1-ER cells obtained from RNA FISH for GATA2 introns and DAPI stain. GATA2 is highly transcribed n this condition.

**Figure S17.**
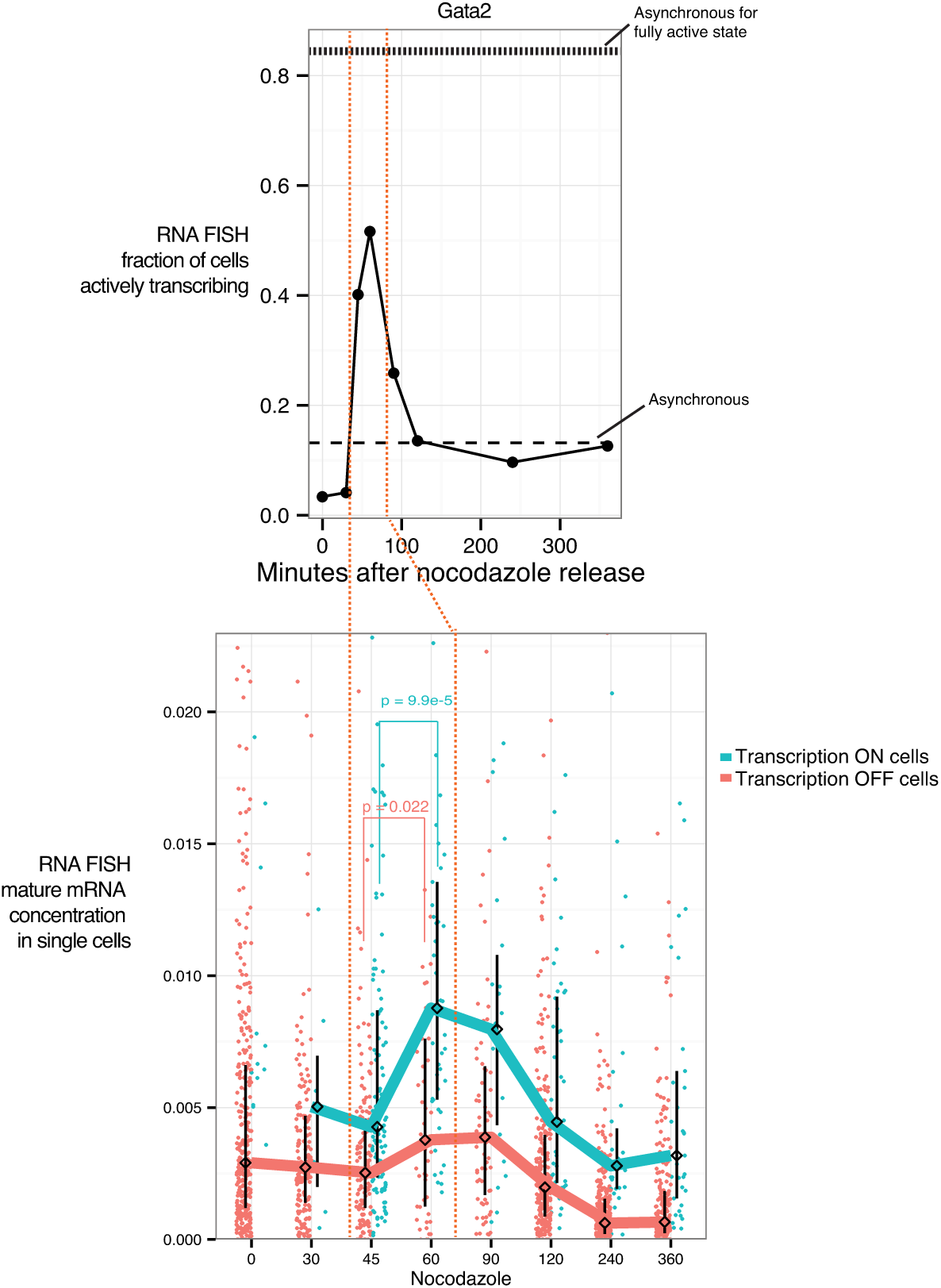
Related to Fig. 5: Additional biological replicates for primary and mature mRNA FISH. We simultaneously imaged Gata2 primary and mature mRNAs by probing for exons and introns in cells synchronized with nocodazole. Shown are data pooled from two biological replicates (combined 89-355 cells for each time point) performed similarly as in Fig. 5. For these replicates, cells were synchronized by nocodazole in the absence of subsequent cell sorting purification, so the 45min-90min time points in particular represent a mixed population of 4N and 2N cells (anaphase to G1). P-values from one-sided Wilcoxon test are shown.

**Figure S18.**
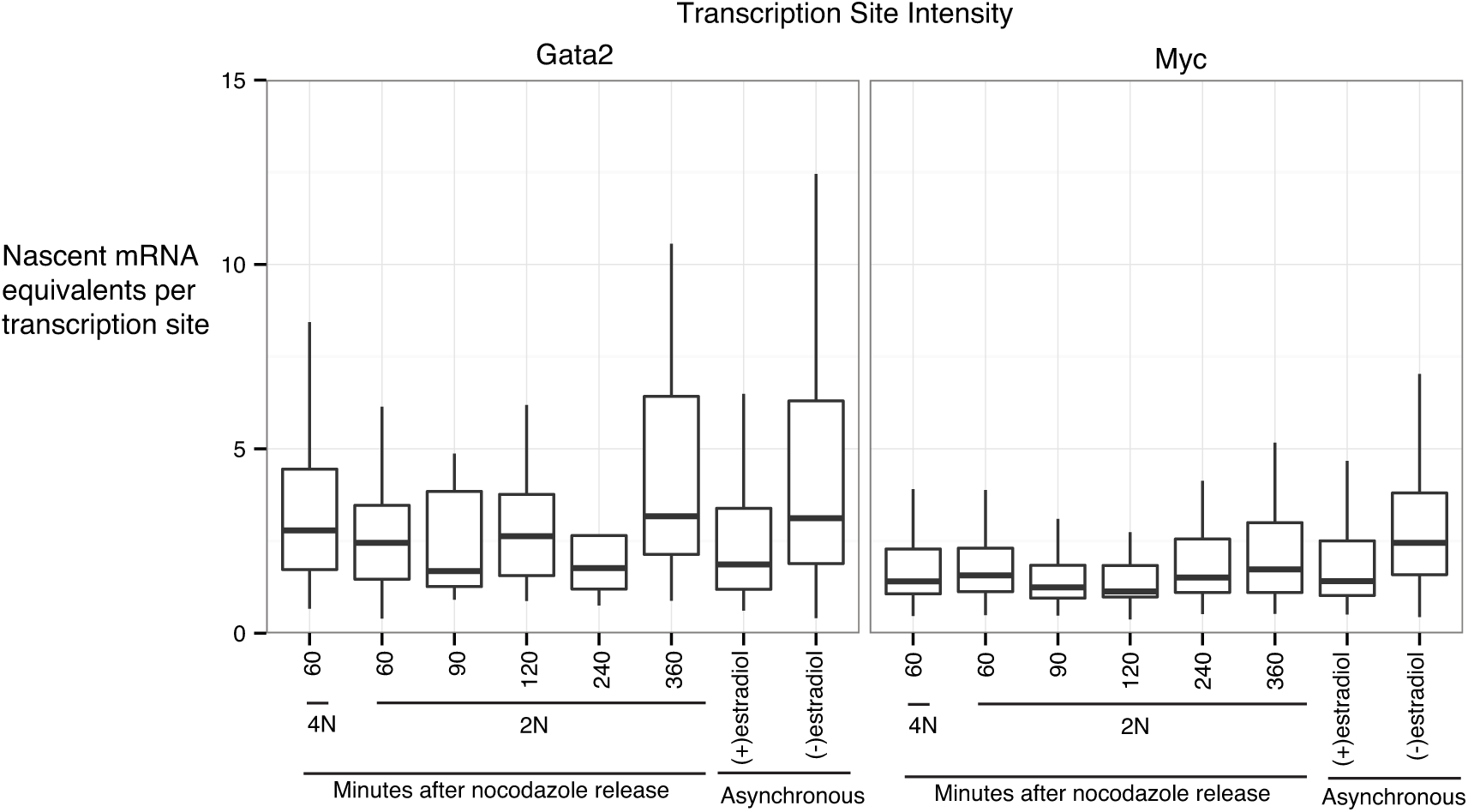
Related to Fig. 5: Transcription site intensities are relatively unchanged with G1 progression. Boxplots of transcription site intensities are shown. Transcription sites are defined as spots where intron and exon spots colocalize, and the intensities are quantified from the exon channel and expressed as equivalents of the average intensity of single mature mRNA molecules.

**Figure S19.**
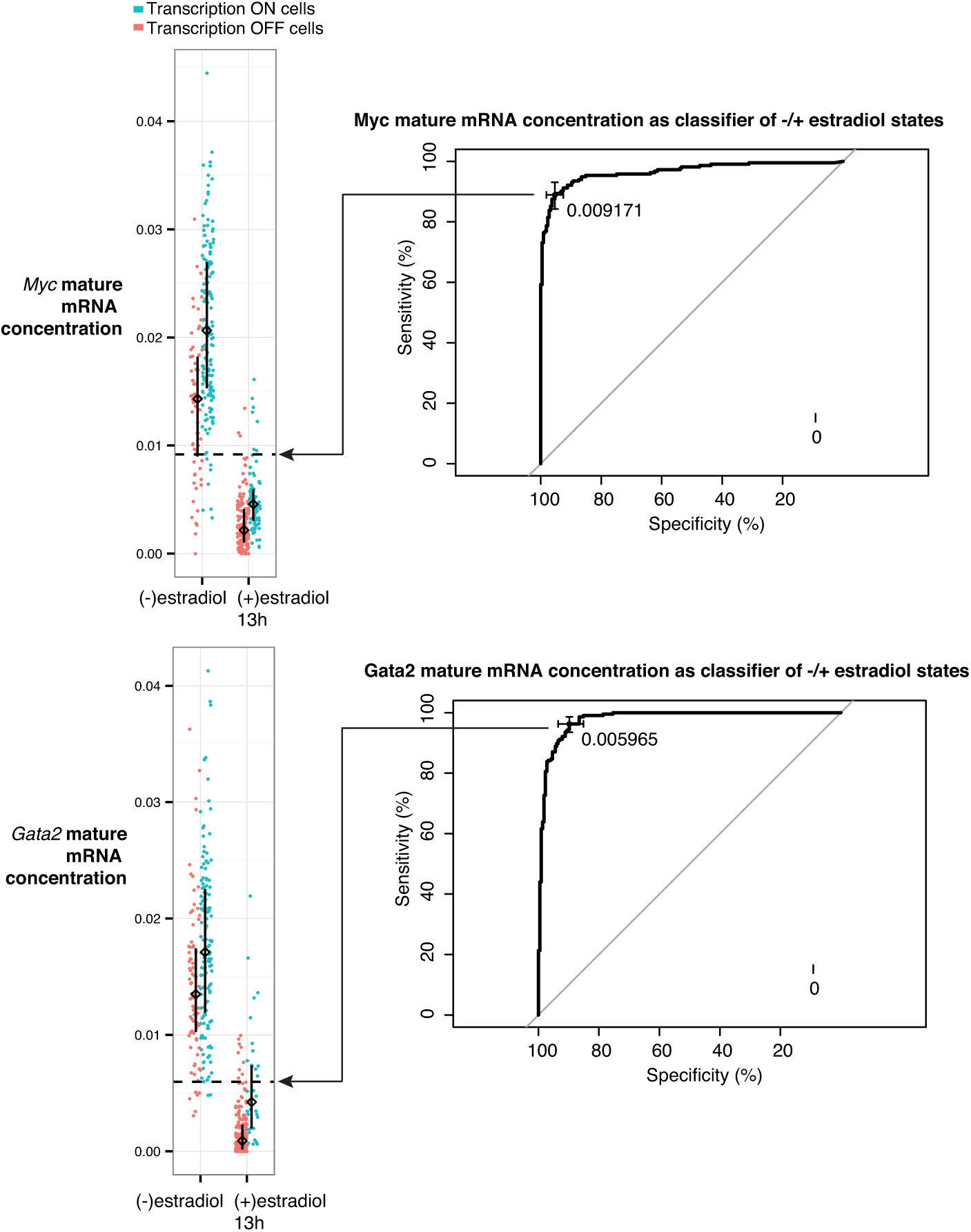
Related to Fig. 5D: Receiver operating characteristics curves for Gata2 and Myc mature mRNA concentrations as classifier for cells in the absence vs. presence of estradiol. Gata2 and Myc mature mRNA concentrations were quantified by RNA FISH for transcriptionally “on” and “off” cells in an asynchronously dividing population of G1E GATA1-ER cells in the absence (−) and presence (+) of estradiol for 13h. The optimal threshold of Gata2 and Myc mature mRNA concentration for discriminating the −/+ estradiol populations was determined by the receiver operating characteristics curves shown above. These the indicated are labeled as “fully active state threshold” in Fig. 5D.

**Figure S20.**
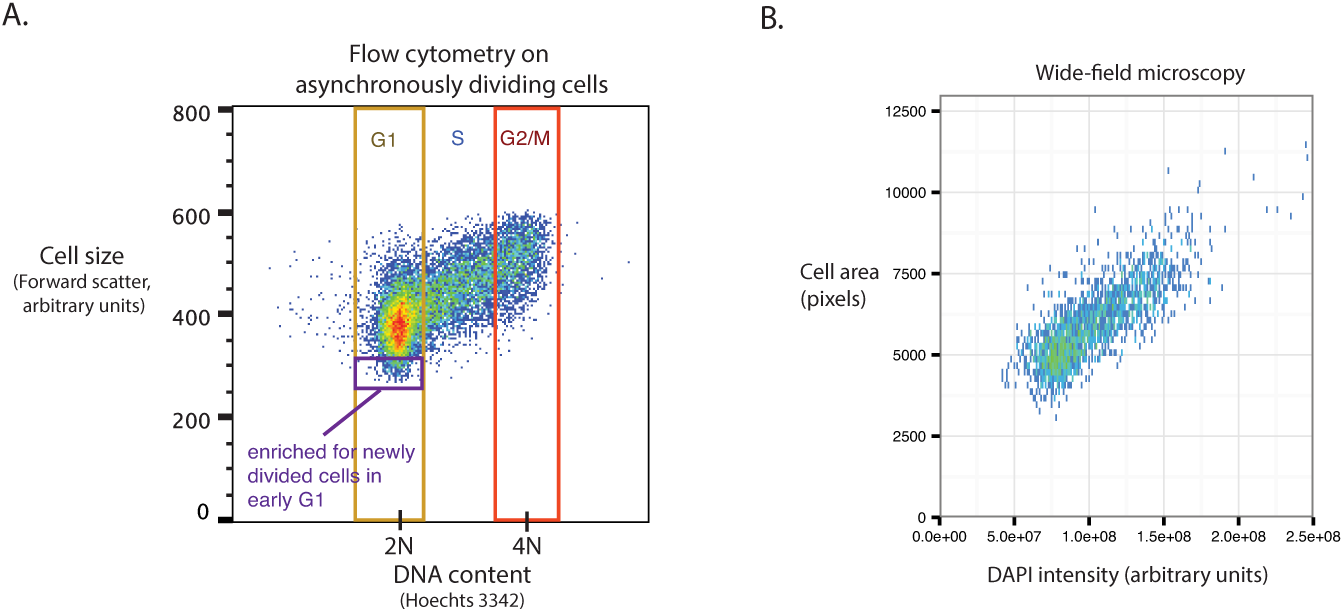
Related to Fig. 5: Cell size is proportional to DNA content in G1E GATA1-ER cells. **A)** Asynchronously dividing G1E GATA1-ER cells were stained with Hoechts 3342 (DNA content) and subjected to flow cytometry. For each cell, forward scatter (reflection of cell size) is plotted against Hoechts 3342, and the boundaries of cell cycle phases are indicate based on DNA content. Within G1, we reason that the smallest cells, highlighted in purple box, are enriched for the newly divided cells in early G1. **B)** We measured cell area (based on manual segmentation of cell boundaries) versus DAPI intensity (sum over single optical plane) by wide-field microscopy, and use their proportionality to estimate thresholds for cell cycle phases in Fig. 6. We further confirmed these cell area thresholds work as approximations of DNA content by arrest-release with nocodazole (analysis not shown).

### Supplemental Experimental Procedures

As described in the Data Access section, we provide scripts that reproduce the majority of figures starting from processed data at an online repository, in addition to raw and processed sequencing data that will be deposited at GEOXXXXXX. We summarize our experimental protocols and computational analyses here.

#### 1 Cell culture, cell cycle synchronization and cell sorting

G1E cells were previously derived through deletion of GATA1 in mouse embryonic stem cells, followed by in vitro differentiation (Weiss et al., 1997). We cultured a sub-line of G1E cells, G1E-ER4, in which GATA1-ER was retrovirally transduced (referred to in main text as “G1E GATA1-ER”), as previously described (Weiss et al., 1997). We retrovirally transduced G1E-ER4 cells with the YFP-MD construct and sorted for a pool of stably YFP-positive cells. Except where indicated in the text as uninduced, we induced cell to mature with 100nM estradiol to activate GATA1-ER. During estradiol induction, we simultaneously treated cells with nocodazole for 7h-13h, washed once, and replated into fresh medium lacking nocodazole for varying times (40min-360min), ensuring all samples are exposed to estradiol for the same duration of 13h.

For ChIP, we harvested cells by resuspension in PBS 2mM EDTA and fixed with 1% formaldehyde at room temperature for 10 minutes with constant mixing, then quenched with 1M glycine, stained with DAPI, and sorted on a BD FACSAria based on YFP-MD and DAPI signal. Sorted fixed cell pellets were snap-frozen with liquid nitrogen and stored at −80 until ready for further processing.

#### 2 Chromatin immunoprecipitation (ChIP)

##### 2.1 Reagent preparation

ChIP-seq of total Pol II was performed in 3 biological replicates using N-20 (Santa Cruz, cat# sc899). For ChIP-qPCR of initiating form of Pol II, we used 8WG16 (Covance, cat# MMS-126R). For H3K27Ac ChIP-seq, we used anti-H3K27Ac from Active-Motif, cat# 39685. For ChIP was performed as follows on approximately 7-20 million cells for each sample. Protease inhibitor (P8340, Sigma) was added to the following buffers right before use: Cell Lysis Buffer (10mM Tris pH 8, 10mM NaCl, 0.2% NP-40/Igpal), Nuclear Lysis Buffer (50mM Tris pH 8, 10mM EDTA, 1% SDS), and IP Dilution Buffer (20mM Tris pH 8, 2mM EDTA, 150mM NaCl, 1% Triton X-100, 0.01% SDS).

Agarose beads slurry was prepared by mixing Protein A (Invitrogen 15918014) and Protein G (Invitrogen 15920010) agarose beads at 1:1 ratio, washed with PBS 3 times, and the mixed beads resuspended in 1:1 volume in PBS (volumes indicated below are for the slurry in PBS, but PBS is removed by centrifugation and aspiration prior to bead use). For use in “pre-clearing” step, 50 µgrabbit IgG was mixed with 50 µL Protein A and G mixed agarose beads slurry. For use for immunoprecipitation step, 70 µL bead slurry was mixed with antibody (20 µg for N20, 10 µg for 8WG1), and 10.2 µg for anti-H3K27Ac for each immunoprecipitation and incubated for >8h at 4^°^C to allow binding (“pre-bound”).

Additional buffers were prepared as follows: IP Wash Buffer 1 (20mM Tris pH 8, 2mM EDTA, 50 mM NaCl, 1% Triton X-100, 0.1% SDS), High Salt Buffer (20 mM Tris pH 8, 2mM EDTA, 500 mM NaCl, 1% Triton X-100, 0.01% SDS), IP Wash Buffer 2 (10 mM Tris pH 8, 1mM EDTA, 0.25 M LiCl, 1% NP-40/Igepal, 1% Na-deoxycholate), Elution Buffer (100 mM NaHCO3, 1% SDS).

##### 2.2 Protocol

All steps performed on ice or at 4^°^C unless otherwise noted. Formaldehyde-fixed cell pellets stored in −80C were thawed on ice, resuspended in 1ml Cell Lysis Buffer and incubated for 20min on ice, then resuspended in 1ml Nuclear Lysis Buffer and incubated for 20min on ice, then diluted with 0.6 ml of IP Dilution Buffer. Samples sonicated at 4^°^C for 45 minutes using either the Bioruptor (Diagenode) or Epishear (Active Motif) (same machine was used within each biological replicate), then cen-trifuged at 21130gfor 10min to remove cellular debris. Supernatant was transferred to new tube and mixed with “pre-clear” beads and rotated at 4^°^C for <5h, then centrifuged at 821g to pellet beads. 200 µL of supernatant was saved as “input.” The remaining supernatant was mixed with beads prebound with antibodies and rotated for > 8h at 4^°^CBeads were washed sequentially once with IP Wash Buffer 1, twice with High Salt Buffer 1, once with IP Wash Buffer 2, then twice with Tris-EDTA pH 8.0 (BP2473-1, Fisher Scientific) (with centrifugation at 5283g for 2min and aspiration of supernatant before resuspension with each subsequent buffer). Then, beads were pelleted by centrifugation and supernatant aspirated.

All steps from this point on performed at room temperature unless otherwise noted. Beads were resuspended in 100 µL Elution Buffer twice sequentially and the supernatant from both steps combined for a final eluate volume of 200 µL. 12 microliter of 5M NaCl and 2 µL of 10mg/ml RNase A (10109169001, BMB) were added to the 200 µL eluted samples and 200 µL input samples and incubated overnight at 65^°^C˙Then,3 µL of 20mg/ml Proteinase K (3115879, BMB) was added and incubation at 65^°^C continued for an additional 2h. 10 µL of 3M sodium acetate pH 5 was added to each sample and DNA purified per the instructions of the QIAquick PCR Purification Kit (cat# 28106, Qiagen). Inputs were eluted in 133.4 µL Buffer EB (Qiagen), and immunoprecipitated samples in 60 µL Buffer EB. DNA was stored at −20^°^C until further processing.

#### 3 ChIP-seq library preparation and Illumina sequencing

All samples, including input, were processed for library construction for Illumina sequencing using Illumina’s TruSeq ChIP Sample Preparation Kit (Illumina cat# IP-202-1012). In brief, DNA fragments were repaired to generate blunt ends, purified using Agencourt AMPure XP Beads (Beckman Coulter cat# A63881), and a single “A” nucleotide was added to each end. Double-stranded Illumina adaptors were ligated to the fragments. Ligation products were purified using Agencourt AMPure XP Beads, and subject to size selection using SPRIselect Beads (Beckman Coulter cat# B23318) in which both a left side size selection was performed at 0.9x volume, and a right side size selection was performed at 0.6x volume according to manufacturer’s specifications. Library fragments were then amplified for 16 cycles of PCR and products were purified using Agencourt AMPure XP Beads. Constructed libraries were run on the Agilent Bioananlyzer 2100 (Agilent Technologies) using either the DNA 7500 kit (cat# 5067-1504) or the High Sensitivity DNA kit (cat# 5067-4626) as appropriate to determine the average size and confirm the absence of unligated adaptors. The mean library size is approximately 330 bp.

The ChIP-seq libraries were quantitated by qPCR using the Kapa SYBR FAST Universal kit (Kapa Biosystems) according to the Illumina’s Sequencing Library qPCR Quantification Guide. Libraries were multiplexed and sequenced on the Illumina HiSeq 2000 using Illumina’s kits and reagents as appropriate.

We sequenced 3 biological replicates for the 0min, 60min, 90min, 180min, and 360min time points; two biological replicates for 240min time point; and one replicate for 40min time point.

#### 4 Bioinformatic analysis of ChIP-seq data

##### 4.1 Read processing

Reads were mapped to mouse mm9 genome using Bowtie. Mapped reads were passed to MACS with a matched control (input) dataset for peak calling for producing bigwig files with reads shifted to account for fragment size. We filtered peaks which had an overlap of at least one base pair with blacklisted regions described in either Pimkin et al. or mm9 blacklisted regions identified by ENCODE. The ENCODE blacklisted regions were obtained from https://sites.google.com/site/anshulkundaje/projects/blacklists. Parameters for the above steps are further detailed in Supplemental File.

##### 4.2 Generation of browser tracks

Bigwig files output by MACS were loaded into Integrated Genomics Viewer. Y-axes were adjusted to normalize for total number of reads mapped in each library.

##### 4.3 Identification of active genes

Because Pol II binding tends to show the greatest enrichment near the 5’ and 3’ ends of genes, we focused on quantification within the 5’ 2.5kb (500bp upstream to 2kb downstream relative to Refseq transcriptional start site) and 3’ 2.5kb (500bp upstream and 2kb downstream relative to Refseq transcriptional end site) regions. If the 5’ or 3’ 2.5kb region of a gene overlapped at least one Pol II peak called by MACS in at least one sample (arrest-release and asynchronous samples with estradiol induction), then we deemed the gene active. From this set of active genes, we removed those whose boundaries (after 500bp extension upstream of transcriptional start site, and 2kb extension downstream of transcriptional end site) overlapped with another active gene. This step is meant to avoid mis-assignment of Pol II signal to the wrong gene. After these steps we arrive at 4309 active and non-overlapping genes that are the subject of Fig. 2 and used for subsequent steps outlined here.

##### 4.4 Identification of intergenic enhancers

We previously predicted distal cis-regulatory modules, or likely enhancers, in G1E ER4 cells based largely on presence of DNase sensitivity and H3K4me1 and relative absence of H3K4me3 (Hsiung et al., 2014). Among these predicted enhancers, we additionally filtered for intergenic enhancers, defined as those that are >20kb away from Refseq gene boundaries and overlap at least one Pol II peak called by MACS in at least one sample (arrest-release and asynchronous samples with estradiol induction). This step is intended to avoid mis-assignment of Pol II signal from binding at genes to enhancers. After these steps we arrive at 809 intergenic enhancers that are the subject of Fig. 3 and used for subsequent steps outlined here.

##### 4.5 Quantitation of Pol II binding

We used bigWigAverageOverBed to count mapped reads and calculated read count per kilobase per million mapped reads in library (RPKM). Changes in RPKM reflect absolute changes in Pol II binding very well even in the context of global changes in Pol II binding Fig. S3, so we use RPKM for all subsequent analyses. For individual replicates and for the mean across replicates, we calculated RPKM at the 5’ and 3’ 2.5kb regions of genes and intergenic enhancers and used them for all analyses that refer to “Pol II binding” in the main text and figures.

##### 4.6 Principal component analysis

We performed principal component analysis on Pol II binding at the 5’ 2.5kb regions of genes, and separately intergenic enhancers, as follows. We confined the principal component analysis to G1 time points (60min, 90min, 180min, 240min, and 360min) in order to focus the analysis on the part of the time course with the patterns of interest. We normalized RPKM of 5’ 2.5kb regions of genes by the sum of RPKM across G1 time points. On these normalized values, we performed principal component analysis using the R package prcomp, with variables scaled to have unit variance prior to analysis (scale. = TRUE). Principal components (“rotation” of the output from prcomp) are plotted against the time points. Projection onto principal components (“x” of the output from prcomp) are referred to as the “degree of match to principal component” in text and figures. The above was performed for individual replicates and for the mean RPKM across replicates. For assessment of replicate concordance in Fig. S6 and Fig. S11, RPKM from individual replicates were projected onto the principal components derived from mean RPKM.

Heatmaps in Fig. 2B and Fig. 3A were generated using the heatmap.2 function from the R package gplots. Rows are median-normalized. The thresholds for separating “early spike” from “late plateau” in Fig. 2B and Fig. 3A were chosen based on the inflection of projection onto the first principal component from positive to negative. The thresholds for separating “late plateau” and “late up-regulated” were chosen manually based on the appearance of the heatmaps.

##### 4.7 Analysis of enriched gene sets

We used GeneTrail (Backes et al., 2007; Keller et al., 2008) to query gene sets from the Gene Ontology, KEGG, and Pfam databases for overrepresentation among the extreme 200 genes based on their projection onto the first principal component for 5’ 2.5kb regions of genes (top and bottom of heatmaps in Fig. 2B). The background is the set of all 4309 active, non-overlapping genes.

##### 4.8 Analysis of chromatin features for association with early G1 transcriptional spike

Below describes how we obtained the Pearson correlation coefficients shown in Fig. 4.

For ChIP-seq signals of H3K27Ac, H3K4me3, H4K4me1, H3K27me3, and H3K9me3, we quantified the read densities of those features within known DNase “hotspots”, which are ¿250bp regions of enriched DNase sensitivity previously categorized as a promoter or distal enhancer hotspot (Hsiung et al., 2014). For 4309 such gene-promoter hotspot pairs, we calculated the Pearson correlation coefficient between the log2 read densities of each histone modification within the promoter DNase hotspot with that of Pol II ChIP-seq projection onto the first principal component for the associated 5’ 2.5kb gene region. For intergenic enhancers, we calculated the Pearson correlation coefficient between the read densities of each histone modification within the intergenic enhancer DNase hotspot with that of Pol II ChIP-seq projection onto the first principal component for those intergenic enhancers.

For DNase-seq signal in mitotic and asynchronous populations, we quantified the read densities of those features within known DNase “peaks”, which are 150bp regions of DNase hypersensitivity (contained within DNase hotspots), previously categorized as residing at promoters or distal enhancers (Hsiung et al., 2014). We calculated the Pearson correlation coefficient between the read densities of each of these features within the promoter DNase peak with that of Pol II ChIP-seq projection onto the first principal component for the associated 5’ 2.5kb gene region. For intergenic enhancers, we calculated the Pearson correlation coefficient between the read densities of each feature within the intergenic enhancer DNase peak with that of Pol II ChIP-seq projection onto the first principal component for DNase hotspot containing those DNase peaks. Where there are multiple DNase peaks within an intergenic enhancer DNase hotspot, we used the DNase peak with the maximum signal for the given feature. The same results are obtained if we use the average of signal among those multiple peaks (not shown). We chose to base these analyses on DNase peaks, rather than hotspots, because they tend to be more dynamic between mitosis and interphase (?).

For GATA1 ChIP-seq signal in mitotic and asynchronous populations, we obtained the union of previously defined GATA1 binding peaks in mitosis and asynchronous cells (Kadauke et al., 2012). We quantified the mitotic or asynchronous GATA1 ChIP-seq read densities within these GATA1 peaks. We assigned each of the 5’ 2.5kb regions of the 4309 active genes to the nearest GATA1 peak (must be ¡ 150kb away). Where there are multiple GATA1 peaks overlapping or equidistant from a given 5’ 2.5kb region, we took the average of read densities across those multiple GATA1 peaks. We calculated the Pearson correlation coefficient between the GATA1 ChIP-seq read densities and the projection onto the Pol II first principal component for each 5’ 2.5kb gene region. The same procedure was applied to intergenic enhancers, except we required that the GATA1 peaks must overlap the intergenic enhancer.

We also performed a similar analysis centered on GATA1 ChIP-seq signal within all DNase peaks (rather than GATA1 binding peaks) that overlap with either the 5’ 2.5kb gene regions or the intergenic enhancer regions, thereby allowing for the correlation coefficient to also include DNase-sensitive regions that do not meet the threshold for calling GATA1 binding peaks. We obtained essentially the same results as in Fig. 4 (not shown).

#### 5. Image analysis

We manually segmented boundaries of cells from brightfield images and localized RNA spots using custom software written in MATLAB (Raj and Tyagi, 2010), with subsequent analyses performed in R. The area within segmentation borders is used for cell area. For Fig. 6, we adjusted for minor systematic variations in the distributions of cell area found across imaging sessions by adding a constant to the cell area, such that the median across all biological replicates are equal. Mature mRNA concentrations per cell are quantified by the spot counts in the exon channel, divided by the cell area.

We took two approaches to quantify co-localization of intron exon spots: 1) manual inspection of each spot by eye to identify relatively bright spots that show co-localization and are thus likely transcription sites, and 2) automated co-localization defined as Gaussian-fitted spots less than 2 pixels apart to identify all likely primary transcripts. The manual inspection approach is more selective for the spots marking nascent transcripts emanating from transcriptionally active gene loci, excluding some co-localized spots of dimmer intensity in the single molecule range that correspond to primary transcripts that have diffused away from the transcription site. On the other hand, the automated approach is easier to apply to a large number of spots and is more precisely defined. Since the presence of primary transcripts that have diffused away from transcription sites indicates recent transcription, the two approaches indeed yield very similar results for Fig. 5 and Fig. S17, so we show the results using the manual inspection approach. For Fig. 6, we used the automated approach because it is easier to implement on a large number of cells. In this case we sum the intensities of primary transcript spot in the exon channel for each cell, expressed as equivalents of the average intensity of a single-molecule mature mRNA spot in the exon channel in a field of cells. Primary transcript equivalents are shown both directly and as values normalized to estimated DNA copy number in Fig. 6.

#### 6. Plotting and graphics

We used R (R Core Team, 2013) packages reshape2 (Wickham, 2007), dplyr (Wickham and Francois, 2015), and ggplot2 (Wickham, 2009) to produce nearly all figures, followed by cosmetic adjustments in Adobe Illustrator.

## References

Akoulitchev, S., and D. Reinberg. 1998. “The Molecular Mechanism of Mitotic Inhibition of TFIIH Is Mediated by Phosphorylation of CDK7.” Genes & Development 12 (22). Cold Spring Harbor Lab: 3541–50

ALONSO’, Angel, Beate Breuer, Barbara Steuer, and Jurgen Fischer. 1991. “The F9-EC Cell Line as a Model for the Analysis of Differentiation.” Int. J. [Je’%lIiol. 35: 389–97

Bender, M. A., Michael Bulger, Jennie Close, and Mark Groudine. 2000. “B-Globin Gene Switching and DNase I Sensitivity of the Endogenous B-Globin LocusinMice Do Not Require the Locus Control Region.” Molecular Cell 5: 387–93

Beyrouthy, Maroun J., Karen E. Alexander, Amy Baldwin, Michael L. Whitfield, Hank W. Bass, Dan McGee, and Myra M. Hurt. 2008. “Identification of G1-Regulated Genes in Normally Cycling Human Cells.” PloS One 3 (12). Public Library of Science: e3943.

Blobel, Gerd A., Stephan Kadauke, Eric Wang, Alan W. Lau, Johannes Zuber, Margaret M. Chou, and Christopher R. Vakoc. 2009. “A Reconfigured Pattern of MLL Occupancy within Mitotic Chromatin Promotes Rapid Transcriptional Reactivation Following Mitotic Exit.” Molecular Cell 36 (6). Elsevier Ltd: 970–83

Campbell, Amy E., Chris C-S Hsiung, and Gerd A. Blobel. 2014. “Comparative Analysis of Mitosis-Specific Antibodies for Bulk Purification of Mitotic Populations by Fluorescence-Activated Cell Sorting.” Bio Techniques 56 (2): 90–91 - 93-94

Caravaca, Juan Manuel, Greg Donahue, Justin S. Becker, Ximiao He, Charles Vinson, and Kenneth S. Zaret. 2013. “Bookmarking by Specific and Nonspecific Binding of FoxA1 Pioneer Factor to Mitotic Chromosomes.” Genes & Development 27 (3). Cold Spring Harbor Lab: 251–60

Christova, Rossitza, and Thomas Oelgeschläger. 2001. “Association of Human TFIID–promoter Complexes with Silenced Mitotic Chromatin in Vivo.” Nature Cell Biology 4 (1). Nature Publishing Group: 79–82

Chubb, Jonathan R., Tatjana Trcek, Shailesh M. Shenoy, and Robert H. Singer. 2006. “Transcriptional Pulsing of a Developmental Gene.” Current Biology: CB 16 (10). Elsevier Ltd: 1018–25

Dani, C., J. M. Blanchard, M. Piechaczyk, S. El Sabouty, L. Marty, and P. Jeanteur. 1984. “Extreme Instability of Myc mRNA in Normal and Transformed Human Cells.” Proceedings of the National Academy of Sciences of the United States of America 81 (22). National Academy of Sciences: 7046–50

Davies, James O. J., Jelena M. Telenius, Simon J. McGowan, Nigel A. Roberts, Stephen Taylor, Douglas R. Higgs, and Jim R. Hughes. 2016. “Multiplexed Analysis of Chromosome Conformation at Vastly Improved Sensitivity.” Nature Methods 13 (1): 74–80

Dey, A., J. Ellenberg, A. Farina, A. E. Coleman, T. Maruyama, S. Sciortino, J. Lippincott-Schwartz, and K. Ozato. 2000. “A Bromodomain Protein, MCAP, Associates with Mitotic Chromosomes and Affects G2-to-M Transition.” Molecular and Cellular Biology 20 (17). American Society for Microbiology: 6537–49

Dey, Anup, Akira Nishiyama, Tatiana Karpova, James McNally, and Keiko Ozato. 2009. “Brd4 Marks Select Genes on Mitotic Chromatin and Directs Postmitotic Transcription.” Molecular Biology of the Cell 20 (23): 4899–4909

Dileep, Vishnu, Ferhat Ay, Jiao Sima, Daniel L. Vera, William S. Noble, and David M. Gilbert. 2015. “Topologically-Associating Domains and Their Long-Range Contacts Are Established during Early G1 Coincident with the Establishment of the Replication Timing Program.” Genome Research, May. Cold Spring Harbor Lab, gr.183699.114.

Dogan, Nergiz, Weisheng Wu, Christapher S. Morrissey, Kuan-Bei Chen, Aaron Stonestrom, Maria Long, Cheryl A. Keller, et al. 2015. “Occupancy by Key Transcription Factors Is a More Accurate Predictor of Enhancer Activity than Histone Modifications or Chromatin Accessibility.” Epigenetics & Chromatin 8 (1). BioMed Central Ltd: 16.

Egli, Dieter, Garrett Birkhoff, and Kevin Eggan. 2008. “Mediators of Reprogramming: Transcription Factors and Transitions through Mitosis.” Nature Publishing Group 9 (7). Nature Publishing Group: 505–16

Femino, A. M. 1998. “Visualization of Single RNA Transcripts in Situ.” Science 280 (5363). American Association for the Advancement of Science: 585–90

Fukuoka, Masashi, Ataru Uehara, Katsuya Niki, Shunya Goto, Dai Kato, Takahiko Utsugi, Masaya Ohtsu, and Yasufumi Murakami. 2012. “Identification of Preferentially Reactivated Genes during Early G1 Phase Using Nascent mRNA as an Index of Transcriptional Activity.” Biochemical and Biophysical Research Communications, December. Elsevier Inc., 1–26

Ganier, Olivier, Stéphane Bocquet, Isabelle Peiffer, Vincent Brochard, Philippe Arnaud, Aurore Puy, Alice Jouneau, Robert Feil, Jean-Paul Renard, and Marcel Méchali. 2011. “Synergic Reprogramming of Mammalian Cells by Combined Exposure to Mitotic Xenopus Egg Extracts and Transcription Factors.” Proceedings of the National Academy of Sciences 108 (42). National Acad Sciences: 17331–36

Glotzer, M., A. W. Murray, and M. W. Kirschner. 1991. “Cyclin Is Degraded by the Ubiquitin Pathway.” Nature 349 (6305): 132–38

Golding, Ido, Johan Paulsson, Scott M. Zawilski, and Edward C. Cox. 2005. “Real-Time Kinetics of Gene Activity in Individual Bacteria.” Cell 123 (6): 1025–36

Gottesfeld, J. M., and D. J. Forbes. 1997. “Mitotic Repression of the Transcriptional Machinery.” Trends in Biochemical Sciences 22 (6): 197–202

Grass, Jeffrey A., Meghan E. Boyer, Saumen Pal, Jing Wu, Mitchell J. Weiss, and Emery H. Bresnick. 2003. “GATA-1-Dependent Transcriptional Repression of GATA-2 via Disruption of Positive Autoregulation and Domain-Wide Chromatin Remodeling.” Proceedings of the National Academy of Sciences of the United States of America 100 (15). National Acad Sciences: 8811–16

Halley-Stott, Richard P., Jerome Jullien, Vincent Pasque, and John Gurdon. 2014. “Mitosis Gives a Brief Window of Opportunity for a Change in Gene Transcription.” PLoS Biology 12 (7). Public Library of Science: e1001914.

Herrick, D. J., and J. Ross. 1994. “The Half-Life of c-Myc mRNA in Growing and Serum-Stimulated Cells: Influence of the Coding and 3’ Untranslated Regions and Role of Ribosome Translocation.” Molecular and Cellular Biology 14 (3). American Society for Microbiology(ASM): 2119–28

Holloway, S. L., M. Glotzer, R. W. King, and A. W. Murray. 1993. “Anaphase Is Initiated by Proteolysis rather than by the Inactivation of Maturation-Promoting Factor.” Cell 73 (7): 1393–1402

Hsiung, Chris C-S, Christapher S. Morrissey, Maheshi Udugama, Christopher L. Frank, Cheryl A. Keller, Songjoon Baek, Belinda Giardine, et al. 2014. “Genome Accessibility Is Widely Preserved and Locally Modulated during Mitosis.” Genome Research 25 (2). Cold Spring Harbor Lab: gr.180646. 114–225

Hughes, Jim R., Jan-Fang Cheng, Nicki Ventress, Shyam Prabhakar, Kevin Clark, Eduardo Anguita, Marco De Gobbi, Pieter de Jong, Eddy Rubin, and Douglas R. Higgs. 2005. “Annotation of Cis-Regulatory Elements by Identification, Subclassification, and Functional Assessment of Multispecies Conserved Sequences.” Proceedings of the National Academy of Sciences of the United States of America 102 (28): 9830–35

Hughes, Jim R., Nigel Roberts, Simon McGowan, Deborah Hay, Eleni Giannoulatou, Magnus Lynch, Marco De Gobbi, Stephen Taylor, Richard Gibbons, and Douglas R. Higgs. 2014. “Analysis of Hundreds of Cis-Regulatory Landscapes at High Resolution in a Single, High-Throughput Experiment.” Nature Genetics 46 (2). Nature Publishing Group: 205–12

Jing, Huie, Christopher R. Vakoc, Lei Ying, Sean Mandat, Hongxin Wang, Xingwu Zheng, and Gerd A. Blobel. 2008. “Exchange of GATA Factors Mediates Transitions in Looped Chromatin Organization at a Developmentally Regulated Gene Locus.” Molecular Cell 29 (2). Elsevier Inc.: 232–42

Kadauke, Stephan, Maheshi I. Udugama, Jan M. Pawlicki, Jordan C. Achtman, Deepti P. Jain, Yong Cheng, Ross C. Hardison, and Gerd A. Blobel. 2012. “Tissue-Specific Mitotic Bookmarking by Hematopoietic Transcription Factor GATA1.” Cell 150 (4). Elsevier Inc.: 725–37

Kelly, Theresa K., Tina Branscombe Miranda, Gangning Liang, Benjamin P. Berman, Joy C. Lin, Amos Tanay, and Peter A. Jones. 2010. “H2A.Z Maintenance during Mitosis Reveals Nucleosome Shifting on Mitotically Silenced Genes.” Molecular Cell 39 (6). Elsevier Inc.: 901–11

Kind, Jop, Ludo Pagie, Havva Ortabozkoyun, Shelagh Boyle, Sandra S. de Vries, Hans Janssen, Mario Amendola, Leisha D. Nolen, Wendy A. Bickmore, and Bas van Steensel. 2013. “Single-Cell Dynamics of Genome-Nuclear Lamina Interactions.” Cell 153 (1): 178–92

Kruhlak, M. J., M. J. Hendzel, W. Fischle, N. R. Bertos, S. Hameed, X. J. Yang, E. Verdin, and D. P. Bazett-Jones. 2001. “Regulation of Global Acetylation in Mitosis through Loss of Histone Acetyltransferases and Deacetylases from Chromatin.” The Journal of Biological Chemistry 276 (41): 38307–19

Kuo, M. Tien, Bhanumathi Iyer, and Robert J. Schwarz. 1982. “Condensation of Chromatin into Chromosomes Preserves an Open Configuration but Alters the DNase I Hypersensitive Cleavage Sites of the Transcribed Gene.” Nucleic Acids Research 10 (15): 4565–79

Lake, Robert J., Pei-Fang Tsai, Inchan Choi, Kyoung-Jae Won, and Hua-Ying Fan. 2014. “RBPJ, the Major Transcriptional Effector of Notch Signaling, Remains Associated with Chromatin throughout Mitosis, Suggesting a Role in Mitotic Bookmarking.” PLoS Genetics 10 (3). Public Library of Science: e1004204.

Langmead, B., C. Trapnell, M. Pop, and S. L. Salzberg. 2009. “Ultrafast and Memory-Efficient Alignment of Short DNA Sequences to the Human Genome.” Genome Biology, January. http://www.biomedcentral.com/content/pdf/gb-2009-10-3-r25.pd.

Lee, Kiwon, Chris C-S Hsiung, Peng Huang, Arjun Raj, and Gerd A. Blobel. 2015. “Dynamic Enhancer-Gene Body Contacts during Transcription Elongation.” Genes & Development 29 (19): 1992–97

Levesque, Marshall J., and Arjun Raj. 2013. “Single-Chromosome Transcriptional Profiling Reveals Chromosomal Gene Expression Regulation.” Nature Methods, February. Nature Publishing Group, 1–6

Lodhi, Niraj, Andrew V. Kossenkov, and Alexei V. Tulin. 2014. “Bookmarking Promoters in Mitotic Chromatin: poly(ADP-Ribose)polymerase-1 as an Epigenetic Mark.” Nucleic Acids Research 42 (11): 7028–38

Martínez-Balbás, M. A., A. Dey, S. K. Rabindran, K. Ozato, and C. Wu. 1995. “Displacement of Sequence-Specific Transcription Factors from Mitotic Chromatin.” Cell 83 (1). Elsevier Inc.: 29–38

Michelotti, E. F., S. Sanford, and D. Levens. 1997. “Marking of Active Genes on Mitotic Chromosomes.” Nature 388 (6645). Nature Publishing Group: 895–99

Muramoto, Tetsuya, Iris MUller, Giles Thomas, Andrew Melvin, and Jonathan R. Chubb. 2010. “Methylation of H3K4 Is Required for Inheritance of Active Transcriptional States.” Current Biology: CB 20 (5). Elsevier Ltd: 397–406

Naumova, Natalia, Maxim Imakaev, Geoffrey Fudenberg, Ye Zhan, Bryan R. Lajoie, Leonid A. Mirny, and Job Dekker. 2013. “Organization of the Mitotic Chromosome.” Science 342 (6161). American Association for the Advancement of Science: 948–53

Padovan-Merhar, Olivia, Gautham P. Nair, Andrew G. Biaesch, Andreas Mayer, Steven Scarfone, Shawn W. Foley, Angela R. Wu, L. Stirling Churchman, Abhyudai Singh, and Arjun Raj. 2015. “Single Mammalian Cells Compensate for Differences in Cellular Volume and DNA Copy Number through Independent Global Transcriptional Mechanisms.” Molecular Cell 58 (2): 339–52

Poleshko, Andrey, Katelyn M. Mansfield, Caroline C. Burlingame, Mark D. Andrake, Neil R. Shah, and Richard A. Katz. 2013. “The Human Protein PRR14 Tethers Heterochromatin to the Nuclear Lamina during Interphase and Mitotic Exit.” CellReports 5 (2). The Authors: 292–301

Prasanth, Kannanganattu V., Paula A. Sacco-Bubulya, Supriya G. Prasanth, and David L. Spector. 2003. “Sequential Entry of Components of Gene Expression Machinery into Daughter Nuclei.” Molecular Biology of the Cell 14 (3). Am Soc Cell Biol: 1043–57

Prescott, D. M., and M. A. Bender. 1962. “Synthesis of RNA and Protein during Mitosis in Mammalian Tissue Culture Cells.” Experimental Cell Research 26 (March): 260–68

Raff, J. W., R. Kellum, and B. Alberts. 1994. “The Drosophila GAGA Transcription Factor Is Associated with Specific Regions of Heterochromatin throughout the Cell Cycle.” The EMBO Journal 13 (24). Nature Publishing Group: 5977.

Raj, Arjun, Charles S. Peskin, Daniel Tranchina, Diana Y. Vargas, and Sanjay Tyagi. 2006. “Stochastic mRNA Synthesis in Mammalian Cells.” PLoS Biology 4 (10): e309.

Raj, Arjun, and Sanjay Tyagi. 2010. Single Molecule Tools: Fluorescence Based Approaches, Part A. Vol. 472. Elsevier Inc.

Raj, Arjun, Patrick van den Bogaard, Scott A. Rifkin, Alexander van Oudenaarden, and Sanjay Tyagi. 2008. “Imaging Individual mRNA Molecules Using Multiple Singly Labeled Probes.” Nature Methods 5 (10). Nature Publishing Group: 877–79

Rizkallah, Raed, Karen E. Alexander, and Myra M. Hurt. 2011. “Global Mitotic Phosphorylation of C2H2 Zinc Finger Protein Linker Peptides.” Cell Cycle 10 (19): 3327–36

Rylski, M., J. J. Welch, Y. Y. Chen, D. L. Letting, J. A. Diehl, L. A. Chodosh, G. A. Blobel, and M. J. Weiss. 2003. “GATA-1-Mediated Proliferation Arrest during Erythroid Maturation.” Molecular and Cellular Biology 23 (14). American Society for Microbiology: 5031–42

Sharova, Lioudmila V., Alexei A. Sharov, Timur Nedorezov, Yulan Piao, Nabeebi Shaik, and Minoru S. H. Ko. 2009. “Database for mRNA Half-Life of 19 977 Genes Obtained by DNA Microarray Analysis of Pluripotent and Differentiating Mouse Embryonic Stem Cells.” DNA Research: An International Journal for Rapid Publication of Reports on Genes and Genomes 16 (1). Oxford University Press: 45–58

Shi, Junwei, Warren A. Whyte, Cinthya J. Zepeda-Mendoza, Joseph P. Milazzo, Chen Shen, Jae-Seok Roe, Jessica L. Minder, et al. 2013. “Role of SWI/SNF in Acute Leukemia Maintenance and Enhancer-Mediated Myc Regulation.” Genes & Development 27 (24): 2648–62

Singh, Amar M., James Chappell, Robert Trost, Li Lin, Tao Wang, Jie Tang, Hao Wu, Shaying Zhao, Peng Jin, and Stephen Dalton. 2013. “Cell-Cycle Control of Developmentally Regulated Transcription Factors Accounts for Heterogeneity in Human Pluripotent Cells.” Stem Cell Reports 1 (6): 532–44

Skinner, Samuel O., Heng Xu, Sonal Nagarkar-Jaiswal, Pablo R. Freire, Thomas P. Zwaka, and Ido Golding. 2016. “Single-Cell Analysis of Transcription Kinetics across the Cell Cycle.” eLife 5 (January). doi:10.7554/eLife.12175.

Smith, Zachary D., Iftach Nachman, Aviv Regev, and Alexander Meissner. 2010. “Dynamic Single-Cell Imaging of Direct Reprogramming Reveals an Early Specifying Event.” Nature Biotechnology 28 (5). Nature Publishing Group: 521–26

Tie, Feng, Rakhee Banerjee, Carl A. Stratton, Jayashree Prasad-Sinha, Vincent Stepanik, Andrei Zlobin, Manuel O. Diaz, Peter C. Scacheri, and Peter J. Harte. 2009. “CBP-Mediated Acetylation of Histone H3 Lysine 27 Antagonizes Drosophila Polycomb Silencing.” Development 136 (18): 3131–41

Varier, Radhika A., Nikolay S. Outchkourov, Petra de Graaf, Frederik M. A. van Schaik, Henk Jan L. Ensing, Fangwei Wang, Jonathan M. G. Higgins, Geert J. P. L. Kops, and Hth Marc Timmers. 2010. “A Phospho/methyl Switch at Histone H3 Regulates TFIID Association with Mitotic Chromosomes.” The EMBO Journal, October. Nature Publishing Group, 1–12

Voichek, Yoav, Raz Bar-Ziv, and Naama Barkai. 2016. “Expression Homeostasis during DNA Replication.” Science 351 (6277): 1087–90

Walter, J. 2003. “Chromosome Order in HeLa Cells Changes during Mitosis and Early G1, but Is Stably Maintained during Subsequent Interphase Stages.” The Journal of Cell Biology 160 (5). Rockefeller Univ Press: 685–97

Wang, Fangwei, and Jonathan M. G. Higgins. 2012. “Histone Modifications and Mitosis: Countermarks, Landmarks, and Bookmarks,” December. Elsevier Ltd, 1–10

Watson, R. J. 1988. “Expression of the c-Myb and c-Myc Genes Is Regulated Independently in Differentiating Mouse Erythroleukemia Cells by Common Processes of Premature Transcription Arrest and Increased mRNA Turnover.” Molecular and Cellular Biology 8 (9). American Society for Microbiology(ASM): 3938–42

Weiss, M. J., C. Yu, and S. H. Orkin. 1997. “Erythroid-Cell-Specific Properties of Transcription Factor GATA-1 Revealed by Phenotypic Rescue of a Gene-Targeted Cell Line.” Molecular and Cellular Biology 17 (3). American Society for Microbiology: 1642–51

Wu, Weisheng, Yong Cheng, Cheryl A. Keller, Jason Ernst, Swathi Ashok Kumar, Tejaswini Mishra, Christapher Morrissey, et al. 2011. “Dynamics of the Epigenetic Landscape during Erythroid Differentiation after GATA1 Restoration.” Genome Research 21 (10): 1659–71

Yang, Jingping, Elizabeth Sung, Paul G. Donlin-Asp, and Victor G. Corces. 2013. “A Subset of Drosophila Myc Sites Remain Associated with Mitotic Chromosomes Colocalized with Insulator Proteins.” Nature Communications 4 (January). Nature Publishing Group: 1464.

Yang, Z., N. He, and Q. Zhou. 2008. “Brd4 Recruits P-TEFb to Chromosomes at Late Mitosis To Promote G1 Gene Expression and Cell Cycle Progression.” Molecular and Cellular Biology 28 (3). American Society for Microbiology: 967–76

Young, Daniel W., Mohammad Q. Hassan, Jitesh Pratap, Mario Galindo, Sayyed K. Zaidi, Suk-Hee Lee, Xiaoqing Yang, et al. 2007. “Mitotic Occupancy and Lineage-Specific Transcriptional Control of rRNA Genes by Runx2.” Nature 445 (7126). Nature Publishing Group: 442–46

Zaidi, Sayyed K., Daniel W. Young, Shirwin M. Pockwinse, Amjad Javed, Jane B. Lian, Janet L. Stein, Andre J. van Wijnen, and Gary S. Stein. 2003. “Mitotic Partitioning and Selective Reorganization of Tissue-Specific Transcription Factors in Progeny Cells.” Proceedings of the National Academy of Sciences of the United States of America 100 (25). National Acad Sciences: 14852–57

Zhang, Yong, Tao Liu, Clifford A. Meyer, Jérôme Eeckhoute, David S. Johnson, Bradley E. Bernstein, Chad Nussbaum, et al. 2008. “Model-Based Analysis of ChIP-Seq (MACS).” Genome Biology 9 (9). BioMed Central Ltd: R137.

Zhao, Rui, Tetsuya Nakamura, Yu Fu, Zsolt Lazar, and David L. Spector. 2011. “Gene Bookmarking Accelerates the Kinetics of Post-Mitotic Transcriptional Re-Activation.” Nature Cell Biology 13 (11). Nature Publishing Group: 1295–1304

## References

Backes, C., Keller, A., Kuentzer, J., Kneissl, B., Comtesse, N., Elnakady, Y. A., Müller, R., Meese, E. and Lenhof, H.-P. (2007). GeneTrail–advanced gene set enrichment analysis. Nucleic Acids Res. 35, W186–92

Hsiung, C. C.-S., Morrissey, C. S., Udugama, M., Frank, C. L., Keller, C. A., Baek, S., Giardine, B., Crawford, G. E., Sung, M.-H., Hardison, R. C. and Blobel, G. A. (2014). Genome accessibility is widely preserved and locally modulated during mitosis. Genome Res. 25, gr.180646.114–225

Kadauke, S., Udugama, M. I., Pawlicki, J. M., Achtman, J. C., Jain, D. P., Cheng, Y., Hardison, R. C. and Blobel, G. A. (2012). Tissue-specific mitotic bookmarking by hematopoietic transcription factor GATA1. Cell 150, 725–737

Keller, A., Backes, C., Al-Awadhi, M., Gerasch, A., Küntzer, J., Kohlbacher, O., Kauf-mann, M. and Lenhof, H.-P. (2008). GeneTrailExpress: a web-based pipeline for the statistical evaluation of microarray experiments. BMC Bioinformatics 9, 552.

McLean, C. Y., Bristor, D., Hiller, M., Clarke, S. L., Schaar, B. T., Lowe, C. B., Wenger, A. M. and Bejerano, G. (2010). GREAT improves functional interpretation of cis-regulatory regions. Nat. Biotechnol. 28, nbt.1630–9.

R Core Team (2013). R: A Language and Environment for Statistiacal Computing. R Foundation for Statistical Computing Vienna, Austria.

Rahl, P. B., Lin, C. Y., Seila, A. C., Flynn, R. A., McCuine, S., Burge, C. B., Sharp, P. A. and Young, R. A. (2010). c-Myc Regulates Transcriptional Pause Release. Cell 141, 432–445

Raj, A. and Tyagi, S. (2010). Single Molecule Tools: Fluorescence Based Approaches, Part A, vol. 472,. Elsevier Inc.

Weiss, M. J., Yu, C. and Orkin, S. H. (1997). Erythroid-cell-specific properties of transcription factor GATA-1 revealed by phenotypic rescue of a gene-targeted cell line. Mol. Cell. Biol. 17, 1642–1651

Wickham, H. (2007). Reshaping Data with the reshape Package. Journal of Statistical Software 21, 1–20

Wickham, H. (2009). ggplot2: elegant graphics for data analysis. Springer New York.

Wickham, H. and Francois, R. (2015). dplyr: A Grammar of Data Manipulation. R package version 0.4.1.

